# Rubisco packaging and stoichiometric composition of a native β-carboxysome

**DOI:** 10.1101/2024.09.20.614183

**Authors:** Yaqi Sun, Yuewen Sheng, Tao Ni, Xingwu Ge, Joscelyn Sarsby, Philip J. Brownridge, Kang Li, Nathan Hardenbrook, Gregory F. Dykes, Nichola Rockliffe, Claire E. Eyers, Peijun Zhang, Lu-Ning Liu

## Abstract

Carboxysomes are anabolic bacterial microcompartments that play an essential role in carbon fixation in cyanobacteria. This self-assembling proteinaceous organelle encapsulates the key CO_2_-fixing enzymes, Rubisco and carbonic anhydrase, using a polyhedral shell constructed by hundreds of shell protein paralogs. Deciphering the precise arrangement and structural organization of Rubisco enzymes within carboxysomes is crucial for understanding the formation process and overall functionality of carboxysomes. Here, we employed cryo-electron tomography and subtomogram averaging to delineate the three-dimensional packaging of Rubiscos within β-carboxysomes in the freshwater cyanobacterium *Synechococcus elongatus* PCC7942 that were grown under low light. Our results revealed that Rubiscos are arranged in multiple concentric layers parallel to the shell within the β-carboxysome lumen. We also identified the binding of Rubisco with the scaffolding protein CcmM in β-carboxysomes, which is instrumental for Rubisco encapsulation and β-carboxysome assembly. Using QconCAT-based quantitative mass spectrometry, we further determined the absolute stoichiometric composition of the entire β-carboxysome. This study and recent findings on the β-carboxysome structure provide insights into the assembly principles and structural variation of β-carboxysomes, which will aid in the rational design and repurposing of carboxysome nanostructures for diverse bioengineering applications.

## Introduction

Carboxysomes are a family of bacterial microcompartments (BMCs) responsible for carbon fixation in cyanobacteria and some proteobacteria (Kerfeld and Erbilgin, 2015; Liu, 2022). The carboxysome sequesters the key CO_2_-fixing enzyme, ribulose 1,5-bisphosphate carboxylase/oxygenase (Rubisco), and carbonic anhydrase (CA), using a polyhedral protein shell (Figure 1A). The carboxysome shell is constructed from multiple protein paralogs (including hexamers, pentamers, and trimers) through self-assembly and is selectively permeable to specific substrates and products, creating elevated levels of CO_2_ around Rubisco in the carboxysome lumen to enhance CO_2_ fixation (Faulkner et al., 2020; Mahinthichaichan et al., 2018). Their intrinsic architectural properties enable carboxysomes to play a crucial role in the global carbon cycle and primary productivity (Behrenfeld et al., 2001; Faulkner et al., 2017; Kerfeld et al., 2018; Rae et al., 2013; Sun et al., 2022; Sun et al., 2019). Understanding the assembly and organizational principles of carboxysomes is of key importance not only for elucidating carboxysome function and catalytic performance but also for harnessing their potential in synthetic engineering for various biotechnological applications (Liu, 2021; Liu et al., 2021).

**Figure 1.**
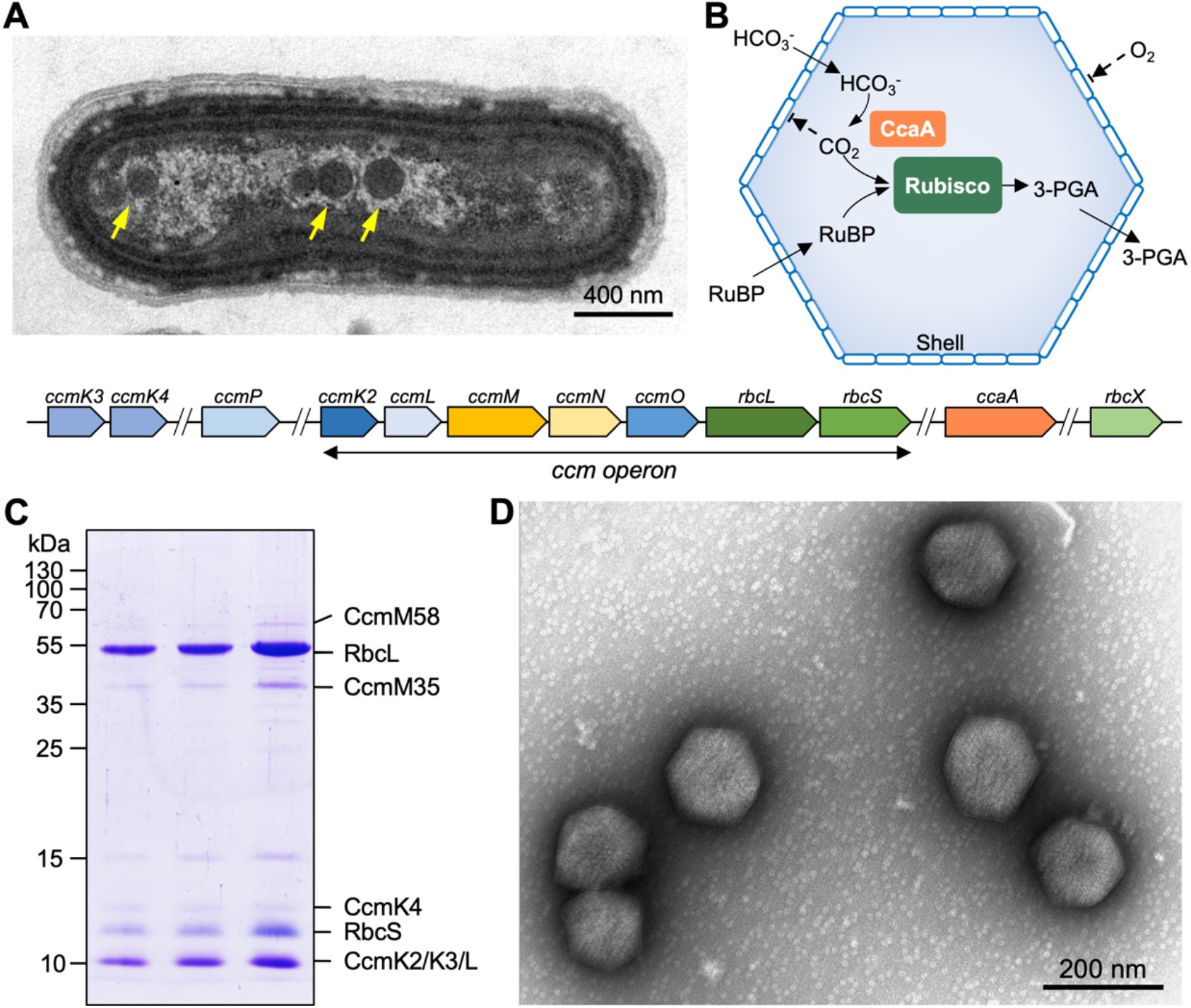
Purification and characterization of Syn7942 β-carboxysomes. (**A**) Thin-section electron microscopy of a Syn7942 cell with β-carboxysomes indicated by yellow arrows (top) and the organization of genes expressing β-carboxysome proteins in the Syn7942 genome (bottom); (**B**) Functional diagrams of β-carboxysomes; (**C**) SDS-PAGE of purified β-carboxysomes from three biological replicates; (**D**) Electron microscopy image of isolated β-carboxysomes from Syn7942.

There are two carboxysome lineages, α-carboxysomes and β-carboxysomes, which differ in the types of encased Rubisco, composition of building proteins, and biogenesis pathway (Kerfeld and Melnicki, 2016; Price et al., 2008; Rae *et al*., 2013). The β-carboxysome of the cyanobacterium *Synechococcus elongatus* PCC7942 (Syn7942) has been extensively characterized as a model carboxysome (Figure 1B). The cargo enzymes of the Syn7942 β-carboxysome include Form IB Rubisco (comprising the large and small subunits RbcL and RbcS, encoded by the *rbcL* and *rbcS* genes respectively) and β-carbonic anhydrase (β-CA, also known as the CcaA protein which is encoded by the *ccaA* gene). The shell consists of the structural proteins CcmK2, CcmK3, and CcmK4, which appear as hexamers and predominantly form shell facets (Kerfeld et al., 2005), CcmL pentamers that occupy the vertices of the polyhedron (Tanaka et al., 2008), as well as the trimeric proteins CcmO and CcmP (Cai et al., 2013; Larsson et al., 2017). In addition, CcmM and CcmN function as “linker” proteins to promote nucleation of Rubisco in the β-carboxysome biogenesis and shell-interior association (Eisenhut et al., 2007; Gonzalez-Esquer et al., 2015; Kinney et al., 2012; Long et al., 2007; Long et al., 2010; Ryan et al., 2019; Wang et al., 2019), which is essential for *de novo* assembly of β-carboxysomes (Cameron et al., 2013; Chen et al., 2013). Moreover, auxiliary factors Raf1 and RbcX are involved in β-carboxysome formation and regulation. Raf1 mediates the assembly of Rubisco holoenzymes and the biogenesis of intact β-carboxysomes (Huang et al., 2020). RbcX serves as a component of the carboxysome and plays a role in mediating β-carboxysome assembly and subcellular distribution (Huang et al., 2019). In the β-carboxysomes that contain CcmK1, a chaperon protein CcmS has recently been determined to play an important role in mediating β-carboxysome formation through interaction with the CcmK1 C-termini exposing at the outer surface of the shell (Chen et al., 2023; Cheng et al., 2024).

Despite the significant efforts made over the past decades to understand the functions of individual carboxysome components, the precise composition of carboxysomes and how the building proteins assemble and self-organize to form intact carboxysome structures remain fundamental questions. Recent studies have provided detailed information on the encapsulation and spatial arrangement of Rubiscos within α-carboxysomes (Evans et al., 2023; Metskas et al., 2022; Ni et al., 2023; Ni et al., 2022b), as well as the exact protein stoichiometry of α-carboxysomes (Sun *et al*., 2022). The β-carboxysomes exhibit structural and compositional variations under different environmental conditions; for example, β-carboxysomes can vary in size, with diameters spanning from 144 to 208 nm, depending on the light and CO_2_ conditions (Sun *et al*., 2019). A recent study characterized the Rubisco organization of β-carboxysomes isolated from Syn7942, which have a diameter of 193 nm (Kong et al., 2024). However, how the Rubisco organization is modulated within the β-carboxysomes of varying sizes, as well as the precise composition of intact β-carboxysomes, remains unclear. This knowledge gap is among the major obstacles in the construction of functional intact β-carboxysomes in non-native organisms for bioengineering applications.

In this study, we islated intact β-carboxysomes with a reduced diameter of 169 nm from Syn7942 cells that were cultivated under low-light conditions and determined the spatial packaging of Rubisco mediated by CcmM within the smaller β-carboxysome using cryo-electron tomography (cryoET). Moreover, we performed absolute quantification of protein components within the native β-carboxysome using QconCAT-assisted quantitative mass spectrometry in combination with biochemical analysis and enzymatic assays. Our results provide structural basis for detailed understanding of the assembly principles and structural regulation of β-carboxysomes. Advanced knowledge of carboxysome assembly and modulation will aid in the rational design and synthetic engineering of carboxysome-based structures.

## Results

### Structure and assembly of Rubisco in isolated Syn7942 β-carboxysomes

Our previous studies indicated that β-carboxysomes produced under low-light conditions tend to be smaller in diameter and more uniform in shape (Sun *et al*., 2019), which may make them suitable for in-depth structural and biochemical analysis. Therefore, we purified intact β-carboxysomes from Syn7942 grown under low light (15 μE·m^−2^·s^−1^) using sucrose gradient ultracentrifugation (Faulkner *et al*., 2017) (Supplementary Figure 1A). The β-carboxysome proteins were identified by sodium dodecyl sulfate–polyacrylamide gel electrophoresis (SDS-PAGE) (Figure 1C) and immunoblot analysis (Supplementary Figure 1B). Negative-staining EM revealed that the isolated β-carboxysomes formed intact polyhedral shapes (Figure 1D). The CO_2_-fixation rate of isolated β-carboxysomes was measured as 2.56 ± 0.16 μmol.mg^−1.^min^−1^ (*n* = 3), which is comparable to that of α-carboxysomes from the chemoautotrophic bacterium *Halothiobacillus neapolitanus* (Sun *et al*., 2022). These results confirmed the functional and structural integrity of β-carboxysomes isolated from Syn7942.

We performed cryoET and subtomogram averaging (STA) of the isolated β-carboxysomes using emClarity (Himes and Zhang, 2018; Ni et al., 2022a) (Figure 2, Supplementary Table 1). The β-carboxysomes exhibited morphological variations with a size of 169.0 ± 11.8 nm (Supplementary Figure 1C), which falls within the range of the diameters determined within and isolated from Syn7942 cells in previous studies (Faulkner *et al*., 2017; Sun *et al*., 2019). Individual Rubiscos inside the β-carboxysomes were readily delineated in the raw tomograms (Figure 2A, Supplementary Figure 2). Template matching and mapping of the position and orientation of individual Rubisco complexes to the original tomograms revealed that Rubiscos had a paracrystalline arrangement within the β-carboxysome lumen, forming four to nine concentric layers (Figure 2B, Supplementary Figure 3A). Among the β-carboxysome particles analyzed, 92.4% had 4-6 layers of Rubiscos, with five layers being the most frequently observed (44.3%, Supplementary Figure 3A). The pairwise distance between neighboring Rubiscos is 121.9 ± 31.4 Å (Supplementary Figure 3B), which is comparable to that of α-carboxysomes (128-129 Å) (Ni *et al*., 2022b). Our analysis reveals that each β-carboxysome contains approximately 639 ± 194 Rubisco complexes (Figure 2C). This quantity is remarkably greater than that of the α-carboxysomes from *Cyanobium* 7001, *H. neapolitanus*, and *Prochlorococcus* (Evans *et al*., 2023; Ni *et al*., 2022b; Zhou et al., 2024). However, the Rubisco content is notably lower than that observed in larger β-carboxysomes with a diameter of 193 nm (Kong *et al*., 2024). This variation in Rubisco content may be attributed mainly to differences in carboxysome diameter, as the spacing between Rubisco pairs remains relatively consistent across these different carboxysomes.

**Figure 2.**
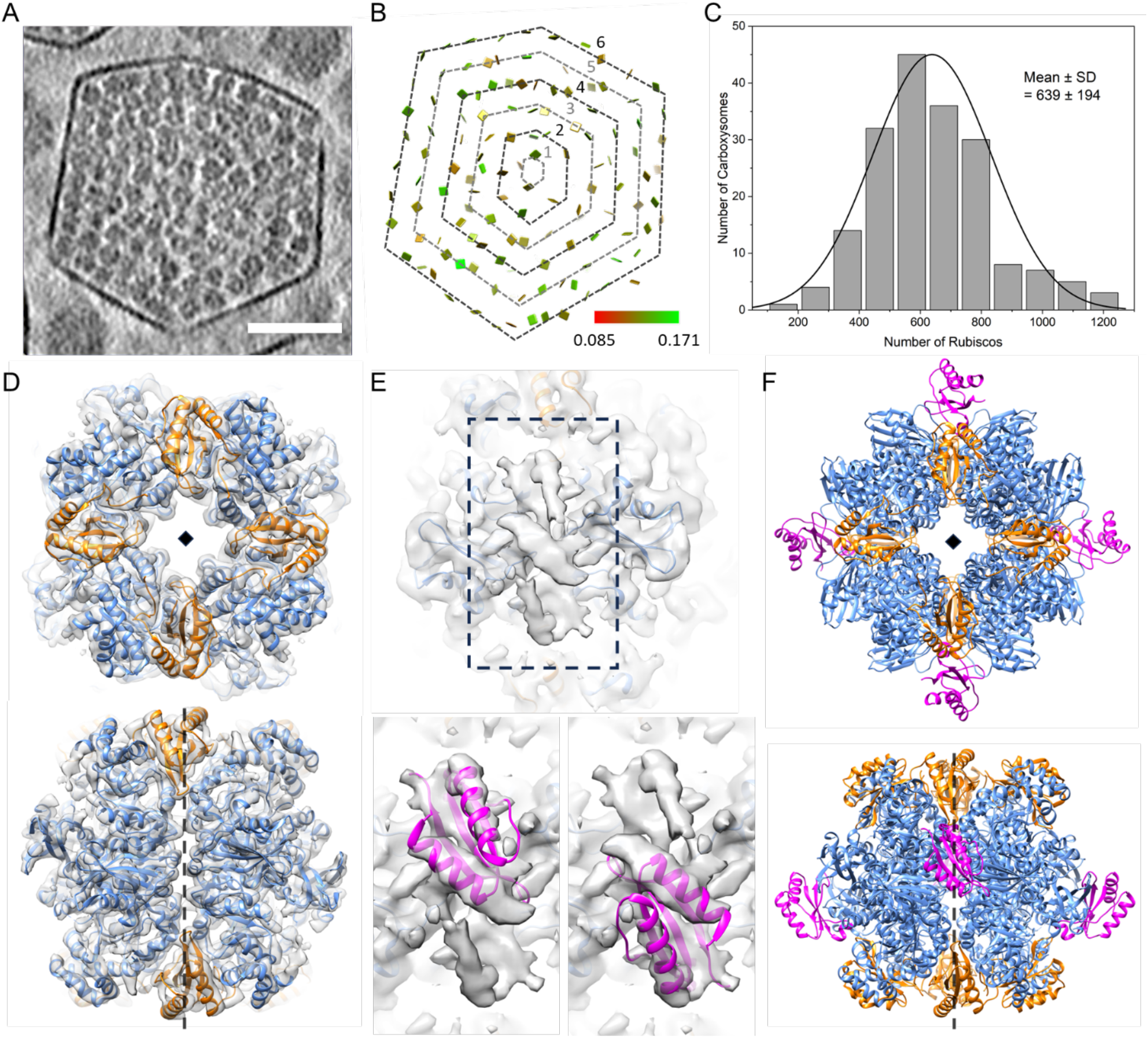
CryoEM structures and organization of Rubisco within β-carboxysomes. (A) A tomogram slice showing a typical beta-carboxysome. Scale bar: 50 nm. (B) The position and orientation of individual rubisco mapped back to the tomogram in (a). It contains a total of 6 layers and the layer number is defined from the core to the shell, as shown. (C) Histogram of Rubisco numbers in β-carboxysomes (*n* = 185). (D) The cryoET subtomogram averaged structures of Rubisco at 3.5 Å, overlapped with refined atomic model (PDB ID: 8BCM), shown in two views. (E) Zoomed-in view showing the density from CcmM, the linker protein which binds Rubisco, and zoomed-in views fitted with atomic model of CcmM SSUL (PDB ID: 6HBC) in two binding modes (upper groove and lower groove). (F) The overall atomic models of Rubisco along with 4 SSUL domains in two views. Only one binding mode is shown (upper groove). The diamond and dashed black lines indicate the fourfold axis.

As expected, the content of Rubisco per layer and total Rubiscos per β-carboxysome gradually increased as the number of layers increased from the core center to the shell (Supplementary Figure 3C-D). Approximately, layers 1-6 contain 1, 20, 71, 174, 308, and 407 copies of Rubiscos, respectively (Supplementary Figure 3C). This leads to approximately 600 copies of Rubiscos per Syn7942 β-carboxysome, given that the majority of β-carboxysomes contain five layers of Rubiscos (Supplementary Figure 3A). The inner four layers are closely packed and relatively ordered; therefore, we only considered the data within the first 2-4 layers to establish a model/formula for the number of Rubiscos for each layer. The prediction was based on the radial distances of Rubiscos from the core as well as the sphere surface area formula (4πr^2^). The average distance between two Rubisco complexes is approximately 120 Å (Supplementary Figure 3B), as is the distance between two consecutive layers. When taking layer 4 as the reference to estimate the total number of Rubiscos for layers 2 and 3, we found that the estimated results closely matched the actual numbers (Supplementary Figure 3C). Accordingly, this allowed us to extrapolate the number of Rubiscos for layers 5-9 in a β-carboxysome that has 9 layers (Supplementary Figure 3D). Moreover, Rubiscos are mostly oriented with their 4-fold axis along the radical direction (90°) across all layers (Supplementary Figure 4A-C). Interestingly, no significant difference in the orientation of Rubisco within the outermost layer compared to the inner layers was observed (Supplementary Figure 4D). This suggests that the size of shell/carboxysome does not have a significant effect on the internal Rubisco arrangement, in line with the “Core first” assembly pathway of β-carboxysomes (Cameron *et al*., 2013). In contrast, a recent study of β-carboxysomes proposed that the Rubiscos at the outermost layer of β-carboxysomes have a more regular arrangement (Kong *et al*., 2024). This discrepancy might be ascribed to the structural plasticity of the β-carboxysomes formed under different growth conditions (see detailed discussion below).

We further determined the structure of Rubisco within native Syn7942 β-carboxysomes at a resolution of 3.5 Å using STA (Figure 2D, Supplementary Figure 2). The resolved structure of the Rubisco L_8_S_8_ hexadecamer was oriented along its 4-fold axis along the radial direction, in good agreement with the reported Syn7942 Rubisco structures (Huang *et al*., 2020; Kong *et al*., 2024; Sun et al., 2024; Wang *et al*., 2019). Intriguingly, an extra density in the groove at the interface between antiparallel RbcL dimers was well-resolved (Figure 2E-2F). This density matches the Rubisco small subunit-like (SSUL) module of the “linker” protein CcmM, which contains two α-helices packed against a β-sheet (Figure 2E). As it sits on the 2-fold axis, the 2-fold symmetrized density appears a half intensity compared to Rubisco, and space can only accommodate one module. We thus deduced that one copy of CcmM SSUL can bind to the RbcL groove in either the up or down configuration, but not both (Figure 2E, Supplementary Figure 5). The binding of CcmM SSUL to Rubisco is reminiscent of that of the reconstituted Rubisco-CcmM35 complex (PDB ID: 6HBC) (Wang *et al*., 2019). Each Rubisco within the carboxysome has potentially four CcmM SSUL domains that can bind Rubisco at the RbcL dimer groove in two different modes (Figure 2E-F). These cryoET results provide direct evidence of Rubisco-CcmM interactions within β-carboxysomes, which drive the recruitment and packaging of Rubisco to form paracrystalline arrays in the β-carboxysome lumen.

### *In situ* cryoET of β-carboxysomes

We further conducted *in situ* cryoET to explore the structure of β-carboxysomes in native Syn7942 cells (Figure 3). The intact β-carboxysome particles exhibit polyhedral shapes and a concentric ring-like arrangement of Rubisco within the carboxysome lumen, corroborating our observations from purified β-carboxysomes (Figure 2). The observed *in vivo* structural variability resembles that observed in previous studies of Syn7942 and the findings of α-carboxysomes (Tsai et al., 2007, Dai et al., 2017, Sun et al., 2022). Additionally, β-carboxysomes were located away from the edge of the cell, often accompanied by large electron-dense granules in proximity (Figure 3). This likely suggests the physical connections between β-carboxysomes and granules, in line with the observations for α-carboxysomes in *H. neapolitanus* cells (Iancu et al., 2010).

**Figure 3.**
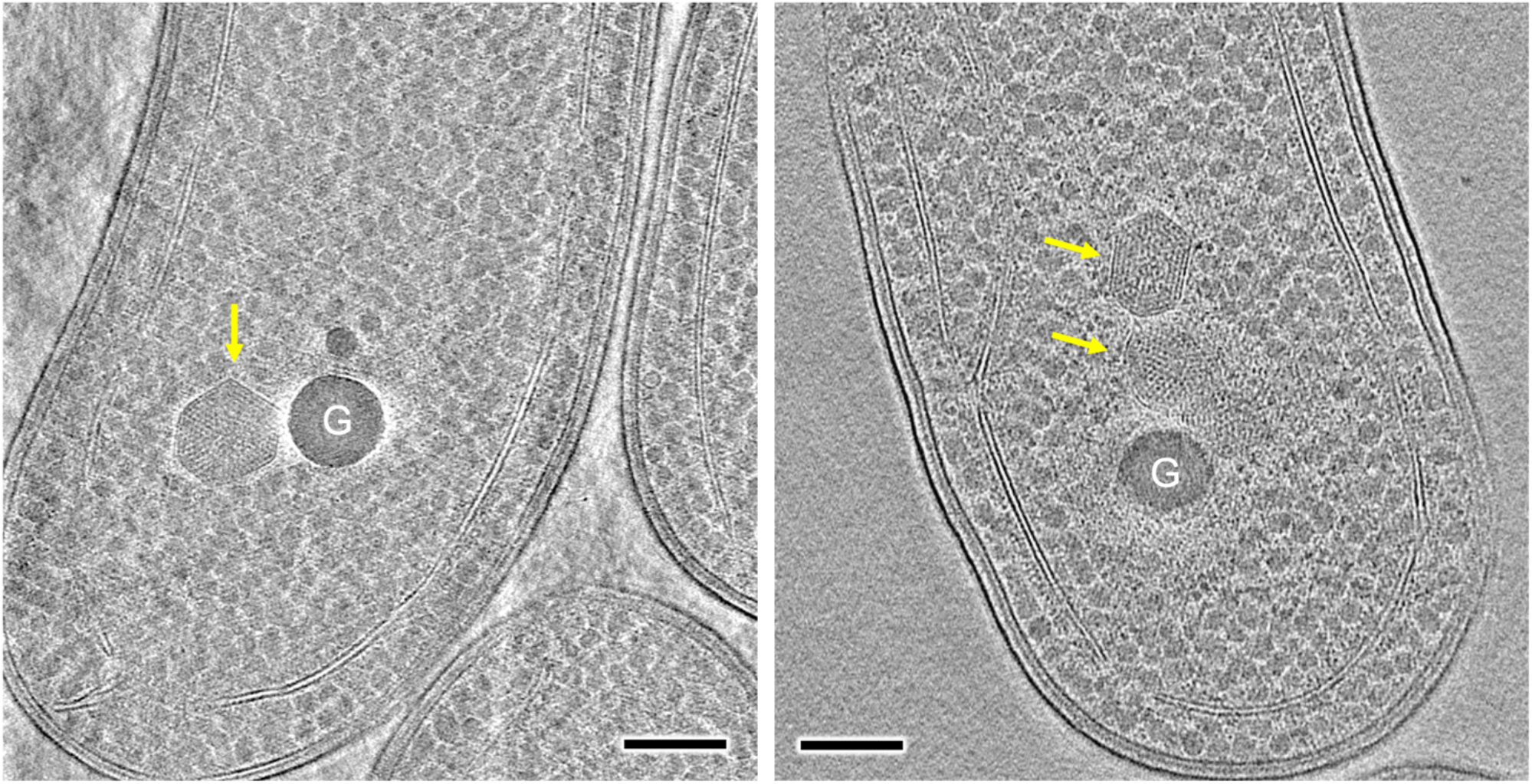
*In situ* cryo-electron tomography (cryoET) of frozen-hydrated Syn7942 cell lamella. Yellow arrows indicate the β-carboxysome structures which possess regularly packed Rubisco in the β-carboxysome lumen. G represents large electron-dense granule. Scale bar, 200 nm.

### Protein stoichiometry of β-carboxysomes

Owing to the irregular morphology of Syn7942 β-carboxysomes, recognition of additional crucial carboxysome components aside from Rubiscos remains unfeasible via the cryoET approach. Using fluorescence tagging and single-molecule fluorescence microscopy, our previous work has estimated the protein abundance of β-carboxysomes *in vivo* (Sun *et al*., 2019). However, given the potential effects of fluorescence tagging and limitations of imaging sensitivity, the precise protein stoichiometry of the β-carboxysome has not been determined. By employing mass spectrometry (MS)-based absolute multiplexed protein quantification using a labelled quantification concatamer (QconCAT) (Johnson et al., 2021; Rivers et al., 2007; Simpson and Beynon, 2012), we previously decoded the accurate stoichiometric composition of protein components of α-carboxysomes and Pdu metabolosomes (Sun *et al*., 2022; Yang et al., 2020). To establish the stoichiometry of the β-carboxysome components, we employed high-resolution liquid chromatography-mass spectrometry (LC-MS) with protein-specific stable isotope-labelled internal standards created using the QconCAT approach. This allowed us to determine the absolute stoichiometry of all the protein components in the purified Syn7942 β-carboxysomes (Figure 4, Table 1). To our knowledge, this is the first precise quantification of the protein stoichiometry of the β-carboxysome.

**Table 1.** Protein stoichiometry of Syn7942 β-carboxysome determined by QconCAT. *Based on the sequence homology of CcmK3 and CcmK4, and of CcmO and CcmP.

| Category | Protein | Structure of functional unit | MW | % of total protein | Units of monomers per carboxysome | Functional units of multimer per carboxysome |
| --- | --- | --- | --- | --- | --- | --- |
| Structural proteins | CcmK2 | Hexamer (PDB: 4OX7) (Cai *et al.*, 2015) | 10.9 kDa | 24.3 ± 2.6 | 11853 ± 2191 | 1975 ± 365 |
| | CcmK3 | Hexamer* | 10.8 kDa | 0.03 ± 0.01 | 12 ± 2 | 2 ± 0 |
| | CcmK4 | Hexamer (PDB: 4OX6) (Cai *et al.*, 2015) | 11.9 kDa | 0.29 ± 0.05 | 127 ± 27 | 21 ± 4 |
| | CcmL | Pentamer (PDB: 2QW7) Tanaka *et al.* 2008) | 11.0 kDa | 0.07 ± 0.02 | 29 ± 8 | 6 ± 2 |
| | CcmO | Pseudo-hexamer* | 29.4 kDa | 0.04 ± 0.05 | 7 ± 7 | 1 ± 1 |
| | CcmP | Pseudo-hexamer (PDB: 5LSR) (Larsson *et al.*, 2017) | 23.2 kDa | 0.14 ± 0.12 | 30 ± 22 | 5 ± 4 |
| | CcmM35 | Monomer | 35.2 kDa | 4.8 ± 1.8 | 730 ± 331 | 730 ± 331 |
| | CcmM58 | Monomer | 57.8 kDa | 1.22 ± 0.21 | 112 ± 27 | 112 ± 27 |
| | CcmN | Monomer | 16.3 kDa | 0.03 ± 0.01 | 8 ± 2 | 8 ± 2 |
| Catalytic proteins | RbcL | L<sub>8</sub>S<sub>8</sub> hexadecamer (PDB: 1RBL) (Newman *et al.*, 1993) | 52.4 kDa | 50.9 ± 4.9 | 5112 | 639 |
| | RbcS | L<sub>8</sub>S<sub>8</sub> hexadecamer (PDB: 1RBL) (Newman *et al.*, 1993) | 13.3 kDa | 17.4 ± 1 | 6919 ± 825 | 865 ± 103 |
| | RbcX | Dimer (PDB: 2PY8) (Tanaka *et al.*, 2007) | 17.0 kDa | 0.03 ± 0.03 | 7 ± 7 | 3 ± 4 |
| | CcaA | Hexamer (PDB: 5SWC) (McGurn *et al.*, 2016) | 30.2 kDa | 0.93 ± 0.76 | 169 ± 151 | 28 ± 25 |
| **Intact β-carboxysome** |  |  | **529.4 MDa** |  |  |  |

**Figure 4.**
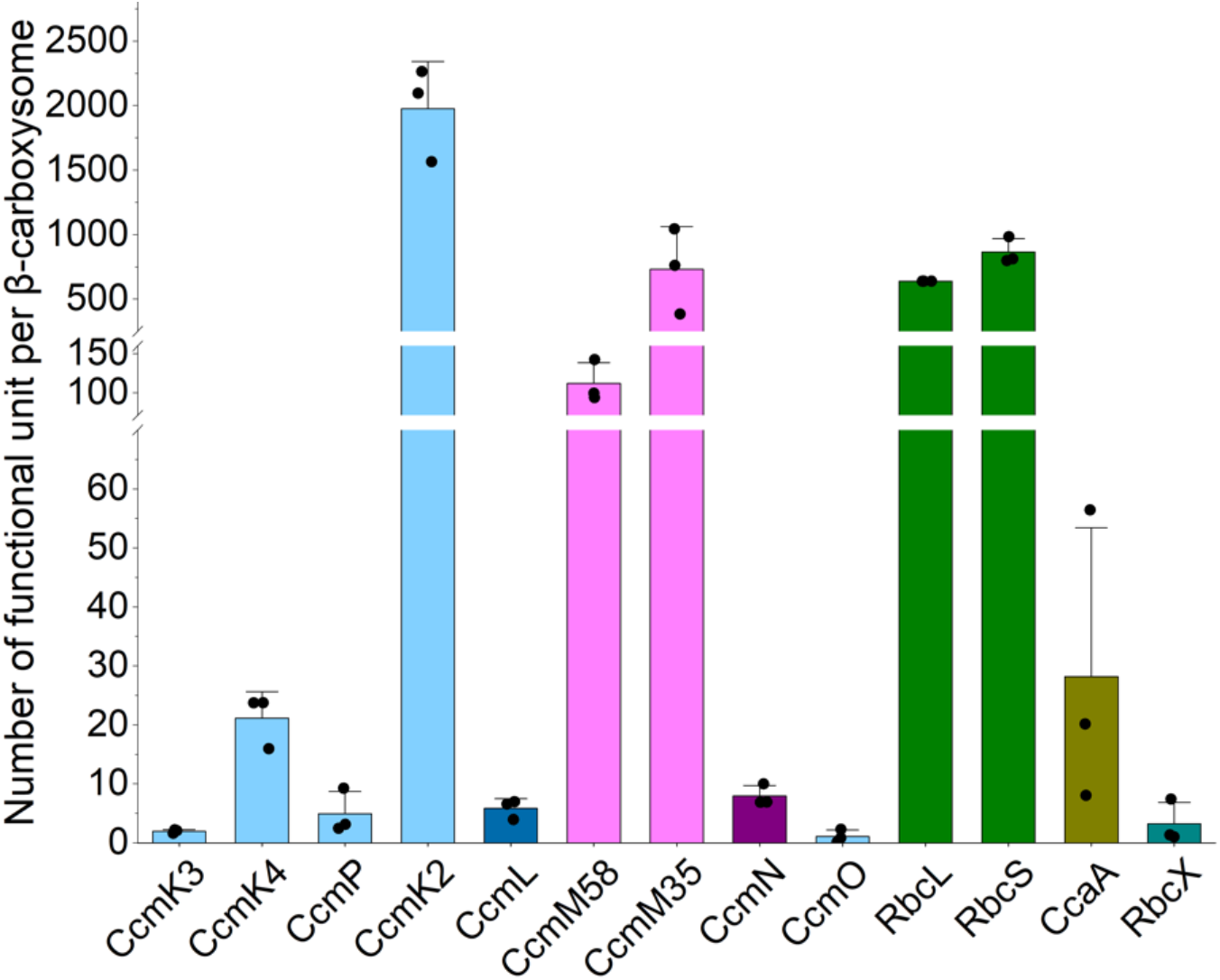
Absolute quantification of the subunit stoichiometry of Syn7942 β-carboxysomes using QconCAT standardization. See also the content of protein oligomers per β-carboxysome in Table 1. Data are presented as means ± SD from four independent biological replicates.

The single QconCAT peptide was designed to contain two unique quantification peptides (Q-peptides) for each Syn7942 β-carboxysome protein: CcmK2, CcmK3, CcmK4, CcmL, the CcmM35/58 shared region, CcmM58, CcmN, CcmO, RbcL, RbcS, CcmP, RbcX, and CcaA (Supplementary Figure 7A, Supplementary Tables 2 and 3). The stable isotope-labelled QconCAT was purified following peptide expression using a cell-free synthesis system (Takemori et al., 2017) (Supplementary Figure 7B), prior to LC-MS-based absolute protein quantification. To determine the absolute amounts of the β-carboxysome protein components, we prepared fresh isolates and co-digested them with purified isotope-labelled QconCAT. All β-carboxysome proteins were successfully detected, enabling us to determine the abundance of protein components within a single β-carboxysome (Figure 4, Table 1). This quantification was based on the Rubisco content determined for the β-carboxysome with an average diameter of 169.0 nm, as measured using cryoET (Supplementary Figure 1C). Our results revealed that the most abundant proteins in the Syn7942 β-carboxysome were CcmK2 hexamers (1975 copies), followed by Rubisco (639 copies, quantified based on the content of RbcL), CcmM35 (730 copies), CcmM58 (112 copies), β-CA (CcaA) hexamers (McGurn et al., 2016) (28 copies), CcmK4 hexamers (21 copies), CcmP pseudo-hexamers (5 copies), CcmN monomers (8 copies), CcmK3 (2 copies of hexamers, see discussion below), CcmO pseudo-hexamers (1 copy), and RbcX dimers (3 copies). Additionally, 6 copies of CcmL pentamers were detected in the β-carboxysome, which is consistent with previously reported observations (Sun *et al*., 2019). Overall, the Syn7942 β-carboxysome had a molecular weight (MW) of ∼529 MDa (Table 1).

In contrast to the low copy numbers of β-CA, Rubisco and CcmK2/K3/K4 hexamers accounted for ∼68% and ∼25% of the total MW, respectively. The Rubisco-scaffolding protein CcmM has two isoforms: a 35-kDa truncated CcmM35 and a full-length 58-kDa CcmM58 (Long *et al*., 2007; Long et al., 2011; Long *et al*., 2010). CcmM35 contains three SSUL domains that interact with Rubisco (Hagen et al., 2018; Wang *et al*., 2019), whereas CcmM58 has an N-terminal γ-CA-like domain that has been suggested to interact with β-CA (Zang et al., 2021), in addition to three C-terminal SSUL domains. Both CcmM35 and CcmM58 accounted for 6% of the total MW. Interestingly, the shell-cargo complex linker protein, CcmN, exhibited a very low abundance, accounting for only 0.02% of the total MW, and was not present in a specific stoichiometric ratio to CcmM58.

## Discussion

Carboxysomes are natural organelle-like microcompartments responsible for carbon fixation in cyanobacteria and some proteobacteria, which serve as important players in the global carbon cycle. Given that they are self-assembling nanostructures composed entirely of proteins, carboxysomes have garnered significant interest for diverse bioengineering applications. In this study, we performed cryoET analysis and absolute quantification using QconCAT-based LC-MS to determine Rubisco packaging and stoichiometric composition of functional, intact Syn7942 β-carboxysomes, a model system widely used in carboxysome studies.

Based on our findings, we propose a model of the entire β-carboxysome structure from Syn7942 (Figure 5). CcmK2 hexamers are the dominant components of the β-carboxysome shell. Other shell proteins, including the CcmK4 and CcmK3 hexamers and the CcmO and CcmP pseudo-hexamers, tile the shell facets. On average, only 6 copies of CcmL pentamers were found per β-carboxysome, which is less than the theoretical estimation of 12 pentagons per canonical icosahedron and the determined number of CsoS4 pentamers (11 copies) per α-carboxysome (Sun *et al*., 2022). This difference likely leads to the greater structural variability of β-carboxysomes compared to that of α-carboxysomes. More than 600 Rubisco L_8_S_8_ hexadecamers were encapsulated within the Syn7942 β-carboxysome with a diameter of 170 nm. Rubiscos are organized in several concentric layers parallel to the outer shell, with Rubisco complexes oriented predominantly in the radial direction (Supplementary Figure 4). The CcmN-CcmM assemblies play an essential role in bridging the association between the shell and cargo enzymes. Our results suggest that there are, on average, 8 copies of CcmN in each β-carboxysome, in contrast to 112 copies of CcmM58 and 730 copies of CcmM35. This is strikingly distinct from the previously estimated CcmN:CcmM stoichiometry based on *in vitro* studies (Sun et al., 2021). Moreover, both our findings and previous structural analysis of reconstituted Rubisco-CcmM complexes (Wang *et al*., 2019) reveal that the SSUL modules of CcmM58 and CcmM35 bind Rubiscos and crosslink neighboring Rubiscos, mediating Rubisco recruitment and packaging within the β-carboxysome. CcmM58 and CcmM35 may crosslink multiple Rubiscos within the same layer or across different layers. Although each Rubisco has four CcmM SSUL-binding sites, our data suggest that the SSUL domains have low occupancy on individual Rubiscos (Figure 2F), which might provide the foundation for the formation of Rubisco condensates and their organizational flexibility. Additionally, there are 28 CcaA enzymes and 3 RbcX proteins encapsulated within the β-carboxysome lumen, which are essential for the assembly and carboxylation activity of Rubiscos.

**Figure 5.**
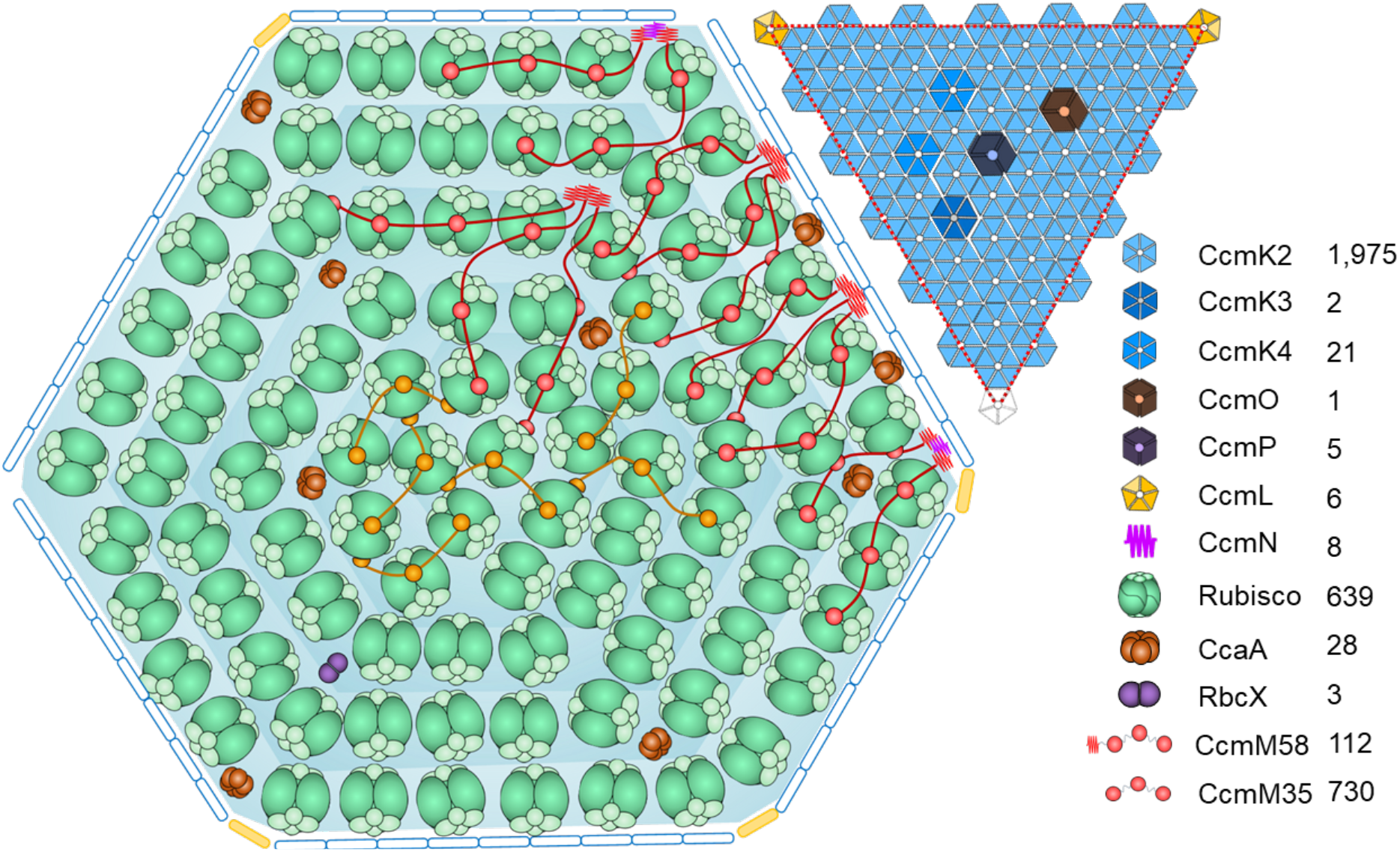
Schematic model of the Syn7942 β-carboxysome structure. The model illustrates the interior organisation and shell facet composition determined by QconCAT-determined stoichiometry and cryoET. Rubiscos (green) are organized in multiple layers in the carboxysome. CcmN (cyan) and the N-terminus of CcmM58 (red fragment) mediate the Rubisco-shell binding. Rubiscos are crosslinked by SSUL of CcmM58 (red) and CcmM35 (orange). CcaA (brown) and RbcX (purple) are co-encapsulated with the Rubisco matrix. On the shell, CcmK3, CcmK4, CcmO, and CcmP are integrated into the facets formed predominantly by CcmK2 hexamers. The vertexes are partially capped by CcmL pentamers.

Our previous studies have revealed a notable size variation of β-carboxysomes in Syn7942 (ranging from 144 to 208 nm) under various environmental conditions, such as light intensity and CO_2_ availability; larger β-carboxysomes can accommodate more Rubisco complexes, with 367 copies of Rubiscos within the β-carboxysome of 144 nm in diameter compared to 1507 Rubiscos within a 208 nm-diameter β-carboxysome (Sun *et al*., 2019). The present work and recent studies (Kong *et al*., 2024) on Syn7942 β-carboxysomes using cryoET further verified this structural variation. Within a β-carboxysome with the diameter of 169 nm, approximately 640 Rubisco enzymes were organized into five concentric layers (Figure 2). A larger β-carboxysome with the size of 193 nm can accommodate an additional Rubisco layer arranged close to the shell, due to the 12 nm distance between neighboring Rubiscos (Supplementary Figure 3), allowing for over 400 additional Rubiscos and more than 1200 Rubisco enzymes in total encapsulated within the larger β-carboxysome structure (Kong *et al*., 2024). The carboxysome diameter, as well as the abundance and packing of interior Rubisco, are important factors in determining the overall architecture and physical mechanics of the intact β-carboxysome (Faulkner *et al*., 2017). Together, these findings underscore the organizational and functional dynamics and flexibility of native β-carboxysomes, a possible mechanism for their adaptation in response to environmental changes.

The findings on β-carboxysomes and α-carboxysomes allow extensive comparisons of the structures and assembly mechanisms of the two distinct carboxysome linkages, although the smaller Syn7942 β-carboxysomes characterized in this study were still larger than the previously reported α-carboxysomes. Recently studies have suggested that the middle region of CsoS2 plays a crucial role in determining the size of the α-carboxysome (Li et al., 2024; Oltrogge et al., 2024). In contrast, what determines the size of β-carboxysomes remains unclear. Our results imply that the composition of scaffolding proteins may drive variations in carboxysome diameter. The ratio of short to long isoforms of CcmM is nearly five times that of CsoS2 (Supplementary Table 4), which presumably favors cargo packaging rather than cargo-shell interactions in the β-carboxysome (Kinney *et al*., 2012; Long *et al*., 2010; Sun *et al*., 2021; Wang *et al*., 2019; Zang *et al*., 2021), resulting in the formation of a large cargo core. In contrast, CcmN, the linker protein that binds directly with shell proteins, has a relatively low abundance (Table 1), suggesting that CcmN is unlikely to be the key factor in determining β-carboxysome size. Nevertheless, despite the difference in the size of the two types of carboxysomes, the ratios between the shell proteins and cargo enzymes remained consistent (Supplementary Table 4), which may be fundamental for maintaining complete encapsulation and overall polyhedral architecture. Although Rubisco packaging and CcmM binding have been identified, how low-abundance proteins, such as CcmN and β-CA, are integrated within the β-carboxysome still remains unclear.

It has been proposed that α- and β-carboxysomes exploit different assembly pathways. *De novo* assembly of β-carboxysomes exploits the “Core first” mode, in which Rubiscos are first condensed, mediated by CcmM, to form a liquid-like Rubisco matrix core, followed by shell encapsulation (Cameron *et al*., 2013; Chen *et al*., 2013). In contrast, the construction of the Rubisco core and shell encapsulation during α-carboxysome assembly, mediated by CsoS2, is assumed to occur in a simultaneous and/or separate fashion (Iancu *et al*., 2010). The cryoET data by us and others revealed that Rubiscos within Syn7942 β-carboxysomes form multiple concentric layers parallel to the shell, which share a similar organizational pattern as Rubiscos in α-carboxysomes from *Cyanobium* and *Prochlorococcus* but are distinct from the Rubisco packaging in *H. neap* α-carboxysomes (Evans *et al*., 2023; Kong *et al*., 2024; Ni *et al*., 2022b; Zhou *et al*., 2024). While the β-carboxysome structures differ in their 3D shape and quantity of concentric layers, our data revealed that there are no significant differences observed in the arrangement and orientation of Rubiscos within each concentric layer among different layers in Syn7942 β-carboxysomes (Figure 2). In addition, the ratios of Rubisco and scaffolding proteins CcmM/CsoS2 in Syn7942 and *H. neap* carboxysomes remained relatively consistent. Overall, the remarkable structural variations in carboxysomes from different native host organisms highlight their modular features and reprogrammable potential, which may be vital for microorganisms to adapt carboxysome functions and CO_2_ assimilation to specific ecological niches (Nguyen et al., 2024; Sun et al., 2016; Sun *et al*., 2019). Our results also suggest that evaluating the architecture of mature carboxysomes alone does not provide a definitive way to differentiate between the distinct assembly mechanisms of α- and β-carboxysomes.

In summary, this study highlights the assembly principles and inherent structural variability of β-carboxysomes, which enables a better understanding of their structural plasticity and functional regulation and facilitates the engineering and reprogramming of carboxysomes for new functions.

## Materials and Methods

### Bacterial strains, growth conditions and carboxysome production

*Synechococcus elongatus* PCC 7942 (Syn7942) cells were cultivated in BG-11 medium (Rippka et al., 1979) or BG-11 agar plates with TES (N-tris(hydroxymethyl)methyl-2-aminoethanesulfonic acid) buffer pH 8.2 (22.9% w/w of C_6_H_15_NO_6_S) and sodium thiosulphate (0.3% w/w of Na_2_S_2_O_3_), solidified on 1.5% agar (w/v). Syn7942 cells were maintained in 5-L Duran flasks under constant low light illumination (15 μE·m^−2^·s^−1^, measured at the culture surface) with aeration and agitation, which enable production of β-carboxysomes with a smaller diameter (Faulkner *et al*., 2017; Sun *et al*., 2019).

### Carboxysome purification from Syn7942

The β-carboxysome purification was performed as previously described with some modifications (Faulkner *et al*., 2017; Sun *et al*., 2016). Syn7942 cells from a 16 L culture were harvested at the late-exponential phase when OD_750_ reached 1.5-2, lysosome-treated, and ruptured by glass beads beating in TE buffer. Lysates were incubated overnight with 1% Triton X-100 and 0.5% IGPAL CA-630 for membrane solubilization. Lysates were briefly centrifuged at 3,000 × g to remove large debris, and the supernatant was centrifuged at 40,000 × g for 1 h to obtain crude β-carboxysome enrichment. The crude enrichments were further treated with 1% DDM (n-dodecyl-β-D-maltoside) and 1% DNase for 2 h and then loaded on an identical step sucrose gradient centrifugation used for recombinant β-carboxysomes. Enriched β-carboxysomes were collected from the 50% to 60% sucrose gradient fractions.

### SDS-PAGE and Immunoblot analysis

SDS-PAGE were performed following standard procedures. 5-10 μg purified carboxysome proteins were loaded per well on 15% polyacrylamide gels and stained with Coomassie Brilliant Blue G-250 (ThermoFisher Scientific, UK). The sizes of the protein bands were referenced to the PageRuler™ Plus Prestained Protein Ladder (ThermoFisher Scientific, UK). Immunoblot analysis was performed using primary rabbit polyclonal anti-RbcL (1:10,000 dilution, AS03 037, Agrisera), rabbit polyclonal anti-CcmK2 (1:5,000 dilution, PHY5336S, Phytoab), working with secondary antibody anti-rabbit IgG secondary antibody (1:10,000 dilution, Agrisera), and horseradish peroxidase-conjugated goat anti-mouse IgG secondary antibody (1:10,000 dilution, Agrisera). Images were acquired using the Quant LAS 4000 platform (GE Healthcare Life Sciences). Quantification of band intensities was performed using ImageJ software.

### Rubisco activity assays

^14^CO_2_ fixation assay was performed to determine the carbon fixation rate of purified carboxysomes, as described previously (Sun *et al*., 2016). Three independently purified carboxysomes were calibrated to 0.2 mg mL^−1^ and added Rubisco assay buffer (100 mm EPPS, pH 8.0, and 20 mm MgCl_2_) at 0.1 μg, assayed was performed at 30°C and initiated with final concentrations of 0.5 mM RuBP. The concentration of HCO_3_ was set to 24 mM for all assays in this study.

### Electron microscopy and data analysis

Electron microscopy was performed as previously described (Chen et al., 2022; Faulkner *et al*., 2017; Huang *et al*., 2020; Sun *et al*., 2022). The purified carboxysomes (∼2 mg mL^−1^) were stained with 3% uranyl acetate on carbon grids and then inspected using an FEI 120 kV Tecnai G2 Spirit BioTWIN TEM equipped with a Gatan Rio 16 camera. The diameters of carboxysomes were measured using ImageJ as described previously (Faulkner *et al*., 2017; Sun *et al*., 2022) and were statistically analyzed using Origin (OriginLab, Massachusetts, USA).

### CryoET sample preparation, data acquisition, and data analysis

To prepare the cryoET grids for β-carboxysomes, 500 µL purified β-carboxysomes in 60% sucrose were diluted to 5 mL with TEMB buffer and concentrated to 100 µL using 100K centrifugal filters (Amicon Ultra) by centrifugation at 700 × g for 10 min to remove sucrose. Then, 100 µL β-carboxysomes were diluted to 5 mL again and concentrated to a final volume of 50 µL (10× concentrated with 0.6% sucrose in TEMB buffer). The concentrated β-carboxysomes were plunge-frozen in ethane onto lacey holy carbon grids (300 mesh, Agar Scientific) using a Leica GP2. The grids were glow-discharged for 45 s before plunge freezing, and gold fiducial beads (7 nm) were mixed with the sample before sample application to the grids. The excess solution was blotted for 3 or 3.5 sec with a humidity of 100% and a temperature of 20°C.

Optimized cryoET grids were loaded onto a Titan Krios microscope (ThermoFisher Scientific) operated at 300 keV in the Electron Bio-Imaging Centre, Diamond). The dataset was acquired using a Gatan Quantum post-column energy filter (Gatan Inc.) operated in zero-loss mode with a 20 eV slit width, paired with a Gatan K3 direct electron detector, using Thermo Scientific Tomography 5 Software by electron counting in super-resolution mode at a physical pixel size of 1.35 per pixel. The tilt series was collected using a dose-symmetric tilt scheme starting from 0° with a 3° tilt increment by a group of three and an angular range of ±60°. The accumulated dose of each tilt series was around 120 e^−^/Å^2^ with a defocus range between −2.5 and −5.5 μm. Ten raw frames were saved for each tilt series. The details of the data collection process are presented in Supplementary Table 1.

For cryoET data analysis, the tilt-series for β-carboxysomes were aligned with IMOD (v4.9.12) (Kremer et al., 1996) using gold fiducials, with the aid of an in-house on-the-fly processing Python script (https://github.com/ffyr2w/cet_toolbox). The center of each identified gold fiducial was checked manually. Subtomogram averaging was performed using emClarity (v1.5.0.4, v1.6.2) (Ni *et al*., 2022a). The Rubisco crystal structure (PDB ID: 5NV3) was converted to a density map at 20 Å resolution with a molmap command in Chimera, and then used as the template for particle searching using emClarity. Template matching was performed using 4× binned tomograms with a pixel size of 5.4 Å (hereafter bin4 tomograms). Rubisco coordinates were manually checked using IMOD to remove false positives and individual Rubiscos outside the β-carboxysome. A total of 185 β-carboxysomes were used for subtomogram averaging and alignment. The averaging and alignment were first performed at bin3 with a pixel size of 4.05 Å for 4 cycles, followed by bin2 with a pixel size of 2.70 Å pixel for 8 cycles, and bin1 with a pixel size of 1.35 Å for 4 cycles. The dataset was divided into two independent subsets during alignment for gold-standard metrics, and the two subsets were combined in the final iteration, resulting in a final resolution of 3.7 Å. D4 symmetry was applied throughout the alignment procedure and the final density map was reconstructed using 2D tilt-series images with cisTEM within the emClarity package at a resolution of 3.5 Å. As for the radial and angular distribution of Rubiscos, the core of each β-carboxysomes was defined as the average of all Rubiscos positions within the β-carboxysome. The distance between each refined Rubisco and the β-carboxysome core was generated to plot the radial distance distribution. A radial vector for each Rubisco was calculated pointing from the core of its corresponding β-carboxysomes to each Rubisco, while the angle between the radial vector and the 4-fold axis or Rubisco was generated to investigate the radial angular distribution.

CryoET of Syn7942 cells was performed as previously described (Huokko et al., 2021). For cell vitrification and cryo-focused ion beam (cryo-FIB) milling, Syn7942 cells were diluted to OD_750_ = ∼0.8 in their culture medium before plunge freezing. 3 µL of suspended cells in culture medium were applied to the front side of glow-discharged R2/2 carbon-coated copper grids (Quantifoil MicroTools), with additional 1 µL cells added to the back side. The grids were blotted for 4 seconds before vitrified in liquid ethane using a Leica GP2 plunge freezer. The vitrified grids were subsequently loaded onto an Aquilos focused ion beam/scanning electron microscope (FIB/SEM) (ThermoFisher Scientific) to prepare thin lamellae of the cells. To reduce overall specimen charging, the grids were sputter-coated with platinum before milling. The ion beam current was gradually adjusted to lower values as the lamellae thinning progress (first using 0.1 nA until 0.75 µm thick, and then 50 pA until ∼ 250 nm thick). For cryoET data collection, tomography tilt-series were acquired from lamella using a Titan Krios microscope (ThermoFisher Scientific) operated at 300 kV. The tilt-series data were acquired with a stage pre-tilt of -7º to account for the angle of lamella, making the lamella surface roughly perpendicular to the electron beam. Tilt-series were collected from -42º to 42º in 2º increment by a group of 3 using dose symmetric scheme (Hagen et al., 2017) in SerialEM (Mastronarde, 2005), with a target defocus of -10 µm. The tilt series reconstruction was performed with a binning factor of 4, resulting in a final pixel size of 2.492 nm. Micrographs were recorded with a K3 camera equipped with a Gatan Quantum energy filter operated in zero-loss mode with a 20 eV slit width. The exposure time was set to 5 seconds and the movies were fractionated into 10 subframes. The nominal magnification of the recorded images was 15000×, with a pixel size of 6.23 Å. The total accumulated dose was ∼120 e^-^/Å^2^.

### Design, cell-free expression and purification of QconCAT standard

Absolute quantification of carboxysome protein components was performed using concatenated signature QconCAT peptides (Pratt et al., 2006) in a manner similar to that described previously (Sun *et al*., 2022; Yang *et al*., 2020). Briefly, two qualified peptide candidates were selected to quantify each protein. Candidate peptides were BLAST searched against the protein database in the Syn7942 database to ensure their uniqueness. The DNA fragment encoding the above peptides, together with GluFib and cMyc, as well as 6x His-tag at the N-terminus and C-terminus, respectively, were generated following the ALACAT/Qbrick assembly strategy, as reported previously (Johnson *et al*., 2021). The final DNA sequence (Table S2) was assembled into a pEU-E01 vector for cell-free expression using wheat germ cell lysates (CellFree Sciences Co. Ltd., Japan). Synthesis was completed with [^13^C_6_, ^15^N_4_] arginine and [^13^C_6_, ^15^N_2_] lysine (CK Isotopes Ltd, UK) using a WEPR8240H full Expression kit following default protocols (2BScientific Ltd, UK). The QconCAT peptides were purified using a HisTrap™ HP Column (Cytiva, UK) following standard methods. The QconCAT was precipitated and resuspended in 30 μL 25 mM ammonium bicarbonate with 0.1% (w/v) RapiGest™ SF surfactant (Waters, UK) and protease inhibitors (Roche cOmplete™, Mini, EDTA-free Protease Inhibitor Cocktail, Merck, UK).

### Proteomic analysis

Proteomic analysis was performed as previously described with modifications (Sun *et al*., 2022). Designed QconCAT Peptides for β-carboxysome protein quantification were listed in Supplementary Table 2. The overall protein and DNA sequence of QconCAT were provided in Supplementary Table 3. For sample handling, the protein concentration of each sample was determined using a NanoDrop Spectrophotometer (ThermoFisher Scientific). Carboxysome samples were diluted to a final protein concentration of 2 μg per 72 μL of 50 mM NH_4_HCO_3_. QconCAT (approximately 0.6 pmol), and the samples were denatured using 8 μL of 10% (w/v) SDS (ThermoFisher Scientific, UK) in 50 mM NH_4_HCO_3_ followed by incubation at 80°C for 10 min. samples were reduced by the addition of 5 μL 12 mM dithiothreitol in 50 mM NH_4_HCO_3_ and incubated at 60°C for 10 min. Alkylation was carried out by adding 5 μL of 36 mM iodoacetamide in 50 mM NH_4_HCO_3_ and incubation at room temperature for 30 min in the dark. treated samples were then bound to 40ng of SP3 Beads (Cytiva, UK) by the addition of acetonitrile to a final concentration of 80% ACN and incubated for 30 min. The beads were washed 3 times with 100% acetonitrile and once with 100% water. The beads were resuspended in 50 mM NH_4_HCO_3_ and the sample was digested with trypsin (1 μL of 200 ng in 50 mM NH_4_HCO_3_ in a final volume of 20 μL). Samples were incubated at 37°C overnight, followed by centrifugation at 17,200 × *g* for 30 min, and transferred to fresh low-binding tubes. Three biological replicates were analyzed using an UltiMate™ 3000 RSLCnano system coupled to a Q Exactive™ HF Hybrid Quadrupole-Orbitrap™ Mass Spectrometer (ThermoFisher Scientific, UK) in data-dependent acquisition mode, according to a previously published protocol (Johnson *et al*., 2021). The LC was operated under the control of a Dionex Chromatography MS Link 2.14. The raw MS data files were loaded into Thermo Proteome Discoverer v.2.4 (ThermoFisher Scientific, UK) and searched against the carboxysome QconCAT database using Mascot v.2.8 (Matrix Science London, UK) with trypsin as the specified enzyme. Each precursor ion was cleanly isolated at the high-resolution scanning speed of the MS1 approach. A precursor mass tolerance of 10 ppm and a fragment ion mass tolerance of 0.01 Da were applied. Data analysis, including run alignment and peak picking, was carried out using Skyline v23.1 (MacCoss Lab Software) (MacLean et al., 2010; Pino et al., 2020). Single carboxysome quantitative normalization was performed using Rubisco counts measured from cryoET, as shown in Figure 2C.

## Data availability

All data needed to evaluate the conclusions in the paper are presented in the paper and/or the supplementary information. The cryoET subtomogram averaging density maps and corresponding atomic models were deposited in the PDB and EMDB, with accession codes 9FWV [https://doi.org/10.2210/pdb9FWV/pdb] and EMD-50836 [https://www.ebi.ac.uk/emdb/EMD-50836], respectively.

## Acknowledgments

We thank the staff at the Liverpool Biomedical Electron Microscopy unit for their support with negative-staining electron microscopy. We thank Diamond Light source for access and support of the cryoEM facilities at the UK National Electron Bio-Imaging Centre (eBIC) (proposal NT21004), funded by the Wellcome Trust, MRC, and BBSRC. Computation was performed at the Diamond Light Source and the Oxford Biomedical Research Computing (BMRC) facility, a joint development between the Wellcome Centre for Human Genetics and the Big Data Institute (BDI) supported by the Health Data Research UK and the NIHR Oxford Biomedical Research Centre. This work was supported by the National Key R&D Program of China (2023YFA0914600, 2021YFA0909600), National Natural Science Foundation of China (32070109), Royal Society (URF\R\180030, RGF\EA\181061), Biotechnology and Biological Sciences Research Council Grant (BB/Y01135X/1, BB/V009729/1, BB/S003339/1), Leverhulme Trust (RPG-2021-286, RPG-2019-300), ERC Advanced Grant (101021133), National Institutes of Health (P50AI150481), and Wellcome Trust Investigator Award (206422/Z/17/Z).

## Author contributions

Yaqi S., P.Z., and L-N.L. conceived the project; Yaqi S. and X.G. performed carboxysome preparation and biochemical characterization; J.S., P.B., Yaqi S., N.R., and C.E.E. conducted mass spectrometry and data analysis; G.F.D. performed negative-staining EM; Yuewen S. and T.N. collected cryoET datasets, processed cryoET data and sub-tomogram averaging; K.L. and N.H. refined Rubisco structures. T.N. performed cryoFIB/SEM lamella preparation, cryoET data collection and reconstruction. Yuewen S., T.N. and P.Z. analyzed structural data. Yaqi S, Yuewen S, P.Z., and L-N.L. wrote the manuscript with contributions from all the authors.

## References

Behrenfeld, M.J., Randerson, J.T., McClain, C.R., Feldman, G.C., Los, S.O., Tucker, C.J., Falkowski, P.G., Field, C.B., Frouin, R., Esaias, W.E., et al. (2001). Biospheric primary production during an ENSO transition. Science 291:2594–2597. 10.1126/science.1055071.

Cai, F., Sutter, M., Bernstein, S.L., Kinney, J.N., and Kerfeld, C.A. (2015). Engineering bacterial microcompartment shells: chimeric shell proteins and chimeric carboxysome shells. ACS Synth Biol 4:444–453. 10.1021/sb500226j.

Cai, F., Sutter, M., Cameron, J.C., Stanley, D.N., Kinney, J.N., and Kerfeld, C.A. (2013). The structure of CcmP, a tandem bacterial microcompartment domain protein from the beta-carboxysome, forms a subcompartment within a microcompartment. J Biol Chem 288:16055–16063. 10.1074/jbc.M113.456897.

Cameron, J.C., Wilson, S.C., Bernstein, S.L., and Kerfeld, C.A. (2013). Biogenesis of a bacterial organelle: the carboxysome assembly pathway. Cell 155:1131–1140. 10.1016/j.cell.2013.10.044.

Chen, A.H., Robinson-Mosher, A., Savage, D.F., Silver, P.A., and Polka, J.K. (2013). The bacterial carbon-fixing organelle is formed by shell envelopment of preassembled cargo. PLoS One 8:e76127. 10.1371/journal.pone.0076127.

Chen, T., Fang, Y., Jiang, Q., Dykes, G.F., Lin, Y., Price, G.D., Long, B.M., and Liu, L.N. (2022). Incorporation of functional Rubisco activases into engineered carboxysomes to enhance carbon fixation. ACS Synth Biol 11:154–161. 10.1021/acssynbio.1c00311.

Chen, X., Zheng, F., Wang, P., and Mi, H. (2023). Novel protein CcmS is required for stabilization of the assembly of β-carboxysome in Synechocystis sp. strain PCC 6803. New Phytol 239:1266–1280. 10.1111/nph.19016.

Cheng, J., Li, C.-Y., Meng, M., Li, J.-X., Liu, S.-J., Cao, H.-Y., Wang, N., Zhang, Y.-Z., and Liu, L.-N (2024). Molecular interactions of the chaperone CcmS and carboxysome shell protein CcmK1 that mediate β-carboxysome assembly. Plant Physiology:kiae438. 10.1093/plphys/kiae438.

Eisenhut, M., Aguirre von Wobeser, E., Jonas, L., Schubert, H., Ibelings, B.W., Bauwe, H., Matthijs, H.C., and Hagemann, M. (2007). Long-term response toward inorganic carbon limitation in wild type and glycolate turnover mutants of the cyanobacterium Synechocystis sp. strain PCC 6803. Plant Physiol 144:1946–1959. 10.1104/pp.107.103341.

Evans, S.L., Al-Hazeem, M.M.J., Mann, D., Smetacek, N., Beavil, A.J., Sun, Y., Chen, T., Dykes, G.F., Liu, L.N., and Bergeron, J.R.C. (2023). Single-particle cryo-EM analysis of the shell architecture and internal organization of an intact α-carboxysome. Structure 31:1–12. 10.1016/j.str.2023.03.008.

Faulkner, M., Rodriguez-Ramos, J., Dykes, G.F., Owen, S.V., Casella, S., Simpson, D.M., Beynon, R.J., and Liu, L.N. (2017). Direct characterization of the native structure and mechanics of cyanobacterial carboxysomes. Nanoscale 9:10662–10673. 10.1039/c7nr02524f.

Faulkner, M., Szabó, I., Weetman, S.L., Sicard, F., Huber, R.G., Bond, P.J., Rosta, E., and Liu, L.-N. (2020). Molecular simulations unravel the molecular principles that mediate selective permeability of carboxysome shell protein. Sci Rep 10:17501. 10.1038/s41598-020-74536-5.

Gonzalez-Esquer, C.R., Shubitowski, T.B., and Kerfeld, C.A. (2015). Streamlined construction of the cyanobacterial CO2-fixing organelle via protein domain fusions for use in plant synthetic biology. Plant Cell 27:2637–2644. 10.1105/tpc.15.00329.

Hagen, A.R., Plegaria, J.S., Sloan, N., Ferlez, B., Aussignargues, C., Burton, R., and Kerfeld, C.A. (2018). In vitro assembly of diverse bacterial microcompartment shell architectures. Nano Lett 18:7030–7037. 10.1021/acs.nanolett.8b02991.

Hagen, W., Wan, W., and Briggs, J. (2017). Implementation of a cryo-electron tomography tilt-scheme optimized for high resolution subtomogram averaging. Journal of Structural Biology 197:191–198. 10.1016/j.jsb.2016.06.007.

Himes, B.A., and Zhang, P. (2018). emClarity: software for high-resolution cryo-electron tomography and subtomogram averaging. Nature Methods 15:955–961. 10.1038/s41592-018-0167-z.

Huang, F., Vasieva, O., Sun, Y., Faulkner, M., Dykes, G.F., Zhao, Z., and Liu, L.N. (2019). Roles of RbcX in carboxysome biosynthesis in the cyanobacterium Synechococcus elongatus PCC7942. Plant Physiol 179:184–194. 10.1104/pp.18.01217.

Huang, F., Kong, W., Sun, Y., Chen, T., Dykes, G.F., Jiang, Y.L., and Liu, L.N. (2020). Rubisco accumulation factor 1 (Raf1) plays essential roles in mediating Rubisco assembly and carboxysome biogenesis. Proc Natl Acad Sci USA 117:17418–17428. 10.1073/pnas.2007990117.

Huokko, T., Ni, T., Dykes, G.F., Simpson, D.M., Brownridge, P., Conradi, F.D., Beynon, R.J., Nixon, P.J., Mullineaux, C.W., Zhang, P., et al. (2021). Probing the biogenesis pathway and dynamics of thylakoid membranes. Nat Commun 12:3475. 10.1038/s41467-021-23680-1.

Iancu, C.V., Morris, D.M., Dou, Z., Heinhorst, S., Cannon, G.C., and Jensen, G.J. (2010). Organization, structure, and assembly of alpha-carboxysomes determined by electron cryotomography of intact cells. J Mol Biol 396:105–117. 10.1016/j.jmb.2009.11.019.

Johnson, J., Harman, V.M., Franco, C., Emmott, E., Rockliffe, N., Sun, Y., Liu, L.-N., Takemori, A., Takemori, N., and Beynon, R.J. (2021). Construction of à la carte QconCAT protein standards for multiplexed quantification of user-specified target proteins. BMC Biology 19:195. 10.1186/s12915-021-01135-9.

Kerfeld, C.A., and Erbilgin, O. (2015). Bacterial microcompartments and the modular construction of microbial metabolism. Trends Microbiol 23:22–34. 10.1016/j.tim.2014.10.003.

Kerfeld, C.A., and Melnicki, M.R. (2016). Assembly, function and evolution of cyanobacterial carboxysomes. Curr Opin Plant Biol 31:66–75. 10.1016/j.pbi.2016.03.009.

Kerfeld, C.A., Aussignargues, C., Zarzycki, J., Cai, F., and Sutter, M. (2018). Bacterial microcompartments. Nat Rev Microbiol 16:277–290. 10.1038/nrmicro.2018.10.

Kerfeld, C.A., Sawaya, M.R., Tanaka, S., Nguyen, C.V., Phillips, M., Beeby, M., and Yeates, T.O. (2005). Protein structures forming the shell of primitive bacterial organelles. Science 309:936–938. 10.1126/science.1113397.

Kinney, J.N., Salmeen, A., Cai, F., and Kerfeld, C.A. (2012). Elucidating essential role of conserved carboxysomal protein CcmN reveals common feature of bacterial microcompartment assembly. J Biol Chem 287:17729–17736. 10.1074/jbc.M112.355305.

Kong, W.W., Zhu, Y., Zhao, H.R., Du, K., Zhou, R.Q., Li, B., Yang, F., Hou, P., Huang, X.H., Chen, Y., et al. (2024). Cryo-electron tomography reveals the packaging pattern of RuBisCOs in Synechococcus beta-carboxysome. Structure 10.1016/j.str.2024.05.007.

Kremer, J.R., Mastronarde, D.N., and McIntosh, J.R. (1996). Computer visualization of three-dimensional image data using IMOD. J Struct Biol 116:71–76. 10.1006/jsbi.1996.0013.

Larsson, A.M., Hasse, D., Valegard, K., and Andersson, I. (2017). Crystal structures of β-carboxysome shell protein CcmP: ligand binding correlates with the closed or open central pore. J Exp Bot 68:3857–3867. 10.1093/jxb/erx070.

Li, T., Chen, T., Chang, P., Ge, X., Chriscoli, V., Dykes, G., Wang, Q., and Liu, L.-N. (2024). Uncovering the roles of the scaffolding protein CsoS2 in mediating the assembly and shape of the α-carboxysome shell. bioRxiv:2024.2005.2014.594188. 10.1101/2024.05.14.594188.

Liu, L.N. (2021). Bacterial metabolosomes: new insights into their structure and bioengineering. Microbial Biotechnology 14:88–93. 10.1111/1751-7915.13740.

Liu, L.N. (2022). Advances in the bacterial organelles for CO2 fixation. Trends in Microbiology 30:567–580. 10.1016/j.tim.2021.10.004.

Liu, L.N., Yang, M., Sun, Y., and Yang, J. (2021). Protein stoichiometry, structural plasticity and regulation of bacterial microcompartments. Curr Opin Microbiol 63:133–141. 10.1016/j.mib.2021.07.006.

Long, B.M., Badger, M.R., Whitney, S.M., and Price, G.D. (2007). Analysis of carboxysomes from Synechococcus PCC7942 reveals multiple Rubisco complexes with carboxysomal proteins CcmM and CcaA. J Biol Chem 282:29323–29335. 10.1074/jbc.M703896200.

Long, B.M., Tucker, L., Badger, M.R., and Price, G.D. (2010). Functional cyanobacterial β-carboxysomes have an absolute requirement for both long and short forms of the CcmM protein. Plant Physiol 153:285–293. 10.1104/pp.110.154948.

Long, B.M., Rae, B.D., Badger, M.R., and Price, G.D. (2011). Over-expression of the beta-carboxysomal CcmM protein in Synechococcus PCC7942 reveals a tight co-regulation of carboxysomal carbonic anhydrase (CcaA) and M58 content. Photosynth Res 109:33–45. 10.1007/s11120-011-9659-8.

MacLean, B., Tomazela, D.M., Shulman, N., Chambers, M., Finney, G.L., Frewen, B., Kern, R., Tabb, D.L., Liebler, D.C., and MacCoss, M.J. (2010). Skyline: an open source document editor for creating and analyzing targeted proteomics experiments. Bioinformatics 26:966–968. 10.1093/bioinformatics/btq054.

Mahinthichaichan, P., Morris, D.M., Wang, Y., Jensen, G.J., and Tajkhorshid, E. (2018). Selective permeability of carboxysome shell pores to anionic molecules. J Phys Chem B 122:9110–9118. 10.1021/acs.jpcb.8b06822.

Mastronarde, D. (2005). Automated electron microscope tomography using robust prediction of specimen movements. Journal of Structural Biology 152:36–51. 10.1016/j.jsb.2005.07.007.

McGurn, L.D., Moazami-Goudarzi, M., White, S.A., Suwal, T., Brar, B., Tang, J.Q., Espie, G.S., and Kimber, M.S. (2016). The structure, kinetics and interactions of the beta-carboxysomal beta-carbonic anhydrase, CcaA. Biochem J 473:4559–4572. 10.1042/BCJ20160773.

Metskas, L.A., Ortega, D., Oltrogge, L.M., Blikstad, C., Lovejoy, D.R., Laughlin, T.G., Savage, D.F., and Jensen, G.J. (2022). Rubisco forms a lattice inside α-carboxysomes. Nat Commun 13:4863. 10.1038/s41467-022-32584-7.

Newman, J., Branden, C.I., and Jones, T.A. (1993). Structure determination and refinement of ribulose 1,5-bisphosphate carboxylase/oxygenase from Synechococcus PCC6301. Acta Crystallogr D Biol Crystallogr 49:548–560. 10.1107/s090744499300530x.

Nguyen, N.D., Pulsford, S.B., and Long, B.M. (2024). Unraveling Rubisco packaging within β-carboxysomes. Structure 32:1023–1025. 10.1016/j.str.2024.07.006.

Ni, T., Frosio, T., Mendonca, L., Sheng, Y., Clare, D., Himes, B.A., and Zhang, P. (2022a). High-resolution in situ structure determination by cryo-electron tomography and subtomogram averaging using emClarity. Nat Protoc 17:421–444. 10.1038/s41596-021-00648-5.

Ni, T., Sun, Y., Burn, W., Al-Hazeem, M.M.J., Zhu, Y., Yu, X., Liu, L.N., and Zhang, P. (2022b). Structure and assembly of cargo Rubisco in two native α-carboxysomes. Nat Commun 13:4299. 10.1038/s41467-022-32004-w.

Ni, T., Jiang, Q., Ng, P.C., Shen, J., Dou, H., Zhu, Y., Radecke, J., Dykes, G.F., Huang, F., Liu, L.-N., et al. (2023). Intrinsically disordered CsoS2 acts as a general molecular thread for α-carboxysome shell assembly. Nature Communications 14:5512. 10.1038/s41467-023-41211-y.

Oltrogge, L.M., Chen, A.W., Chaijarasphong, T., Turnšek, J.B., and Savage, D.F. (2024). α-Carboxysome Size Is Controlled by the Disordered Scaffold Protein CsoS2. Biochemistry 63:219–229. 10.1021/acs.biochem.3c00403.

Pino, L.K., Searle, B.C., Bollinger, J.G., Nunn, B., MacLean, B., and MacCoss, M.J. (2020). The Skyline ecosystem: Informatics for quantitative mass spectrometry proteomics. Mass Spectrom Rev 39:229–244. 10.1002/mas.21540.

Pratt, J.M., Simpson, D.M., Doherty, M.K., Rivers, J., Gaskell, S.J., and Beynon, R.J. (2006). Multiplexed absolute quantification for proteomics using concatenated signature peptides encoded by QconCAT genes. Nat Protoc 1:1029–1043. 10.1038/nprot.2006.129.

Price, G.D., Badger, M.R., Woodger, F.J., and Long, B.M. (2008). Advances in understanding the cyanobacterial CO2-concentrating-mechanism (CCM): functional components, Ci transporters, diversity, genetic regulation and prospects for engineering into plants. J Exp Bot 59:1441–1461. 10.1093/jxb/erm112.

Rae, B.D., Long, B.M., Badger, M.R., and Price, G.D. (2013). Functions, compositions, and evolution of the two types of carboxysomes: polyhedral microcompartments that facilitate CO2 fixation in cyanobacteria and some proteobacteria. Microbiol Mol Biol Rev 77:357–379. 10.1128/MMBR.00061-12.

Rippka, R., Deruelles, J., Waterbury, J.B., Herdman, M., and Stanier, R.Y. (1979). Generic Assignments, Strain Histories and Properties of Pure Cultures of Cyanobacteria. Microbiology 111:1–61. 10.1099/00221287-111-1-1.

Rivers, J., Simpson, D.M., Robertson, D.H., Gaskell, S.J., and Beynon, R.J. (2007). Absolute multiplexed quantitative analysis of protein expression during muscle development using QconCAT. Molecular & cellular proteomics : MCP 6:1416–1427. 10.1074/mcp.M600456-MCP200.

Ryan, P., Forrester, T.J.B., Wroblewski, C., Kenney, T.M.G., Kitova, E.N., Klassen, J.S., and Kimber, M.S. (2019). The small RbcS-like domains of the beta-carboxysome structural protein CcmM bind RubisCO at a site distinct from that binding the RbcS subunit. J Biol Chem 294:2593–2603. 10.1074/jbc.RA118.006330.

Simpson, D.M., and Beynon, R.J. (2012). QconCATs: design and expression of concatenated protein standards for multiplexed protein quantification. Analytical and bioanalytical chemistry 404:977–989. 10.1007/s00216-012-6230-1.

Sun, H., Cui, N., Han, S.J., Chen, Z.P., Xia, L.Y., Chen, Y., Jiang, Y.L., and Zhou, C.Z. (2021). Complex structure reveals CcmM and CcmN form a heterotrimeric adaptor in beta-carboxysome. Protein Sci 30:1566–1576. 10.1002/pro.4090.

Sun, Y., Wollman, A.J.M., Huang, F., Leake, M.C., and Liu, L.N. (2019). Single-organelle quantification reveals the stoichiometric and structural variability of carboxysomes dependent on the environment. Plant Cell 31:1648–1664. 10.1105/tpc.18.00787.

Sun, Y., Casella, S., Fang, Y., Huang, F., Faulkner, M., Barrett, S., and Liu, L.N. (2016). Light Modulates the Biosynthesis and Organization of Cyanobacterial Carbon Fixation Machinery through Photosynthetic Electron Flow. Plant Physiol 171:530–541. 10.1104/pp.16.00107.

Sun, Y., Chen, T., Ge, X., Ni, T., Dykes, G.F., Zhang, P., Huang, F., and Liu, L.N. (2024). Engineering functional CO2-fixing modules in E. coli via efficient assembly of cyanobacterial Rubisco and carboxysomes. Research Square 10.21203/rs.3.rs-4511266/v1.

Sun, Y., Harman, V.M., Johnson, J.R., Brownridge, P.J., Chen, T., Dykes, G.F., Lin, Y., Beynon, R.J., and Liu, L.N. (2022). Decoding the absolute stoichiometric composition and structural plasticity of α-carboxysomes. mBio 13:e0362921. 10.1128/mbio.03629-21.

Takemori, N., Takemori, A., Tanaka, Y., Endo, Y., Hurst, J.L., Gomez-Baena, G., Harman, V.M., and Beynon, R.J. (2017). MEERCAT: Multiplexed Efficient Cell Free Expression of Recombinant QconCATs For Large Scale Absolute Proteome Quantification. Mol Cell Proteomics 16:2169–2183. 10.1074/mcp.RA117.000284.

Tanaka, S., Sawaya, M.R., Kerfeld, C.A., and Yeates, T.O. (2007). Structure of the RuBisCO chaperone RbcX from Synechocystis sp. PCC6803. Acta Crystallogr D Biol Crystallogr 63:1109–1112. 10.1107/S090744490704228X.

Tanaka, S., Kerfeld, C.A., Sawaya, M.R., Cai, F., Heinhorst, S., Cannon, G.C., and Yeates, T.O. (2008). Atomic-level models of the bacterial carboxysome shell. Science 319:1083–1086. 10.1126/science.1151458.

Wang, H., Yan, X., Aigner, H., Bracher, A., Nguyen, N.D., Hee, W.Y., Long, B.M., Price, G.D., Hartl, F.U., and Hayer-Hartl, M. (2019). Rubisco condensate formation by CcmM in β-carboxysome biogenesis. Nature 566:131–135. 10.1038/s41586-019-0880-5.

Yang, M., Simpson, D.M., Wenner, N., Brownridge, P., Harman, V.M., Hinton, J.C.D., Beynon, R.J., and Liu, L.N. (2020). Decoding the stoichiometric composition and organisation of bacterial metabolosomes. Nat Commun 11:1976. 10.1038/s41467-020-15888-4.

Zang, K., Wang, H., Hartl, F.U., and Hayer-Hartl, M. (2021). Scaffolding protein CcmM directs multiprotein phase separation in beta-carboxysome biogenesis. Nat Struct Mol Biol 28:909–922. 10.1038/s41594-021-00676-5.

Zhou, R.-Q., Jiang, Y.-L., Li, H., Hou, P., Kong, W.-W., Deng, J.-X., Chen, Y., Zhou, C.-Z., and Zeng, Q. (2024). Structure and assembly of the α-carboxysome in the marine cyanobacterium Prochlorococcus. Nature Plants 10:661–672. 10.1038/s41477-024-01660-9.

