## Supplementary Information for "Rubisco packaging and stoichiometric composition of a native β-carboxysome"

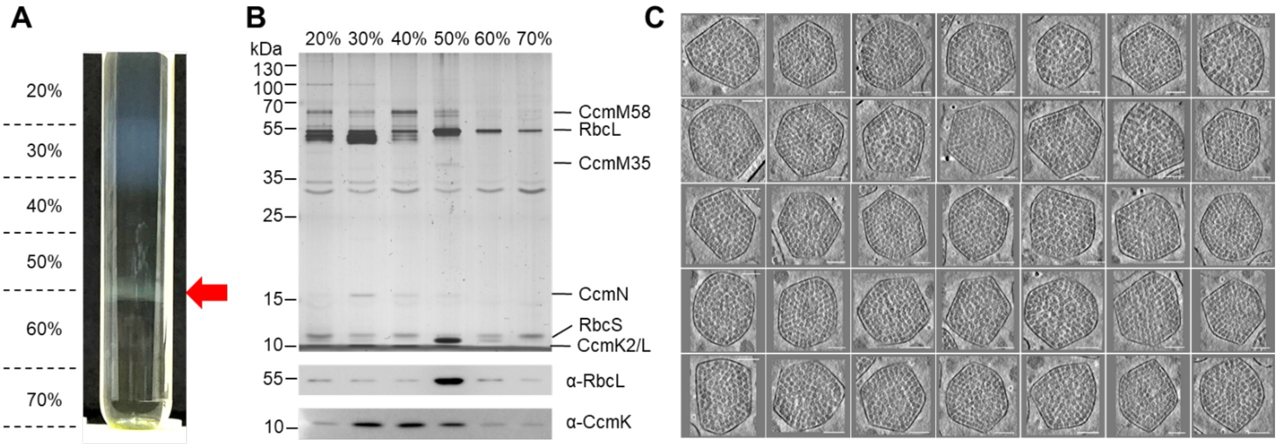


**Supplementary Figure 1. Isolation and identification of Syn7942 β-carboxysomes.** (**A**) Sucrose gradient purification of β-carboxysomes from Syn7942, the red arrow indicates enriched carboxysomes; (**B**) SDS-PAGE and immunoblots identification for Syn7942 β-carboxysomes in sucrose gradient fractions. The major enzyme Rubisco and main shell CcmK2 were detected by RbcL and CcmK2 antibodies respectively. Intact β-carboxysomes are enriched at the bottom of the 50% sucrose gradient; (**C**) Morphological variations of Syn7942 β-carboxysomes. Scale bar: 50 nm.

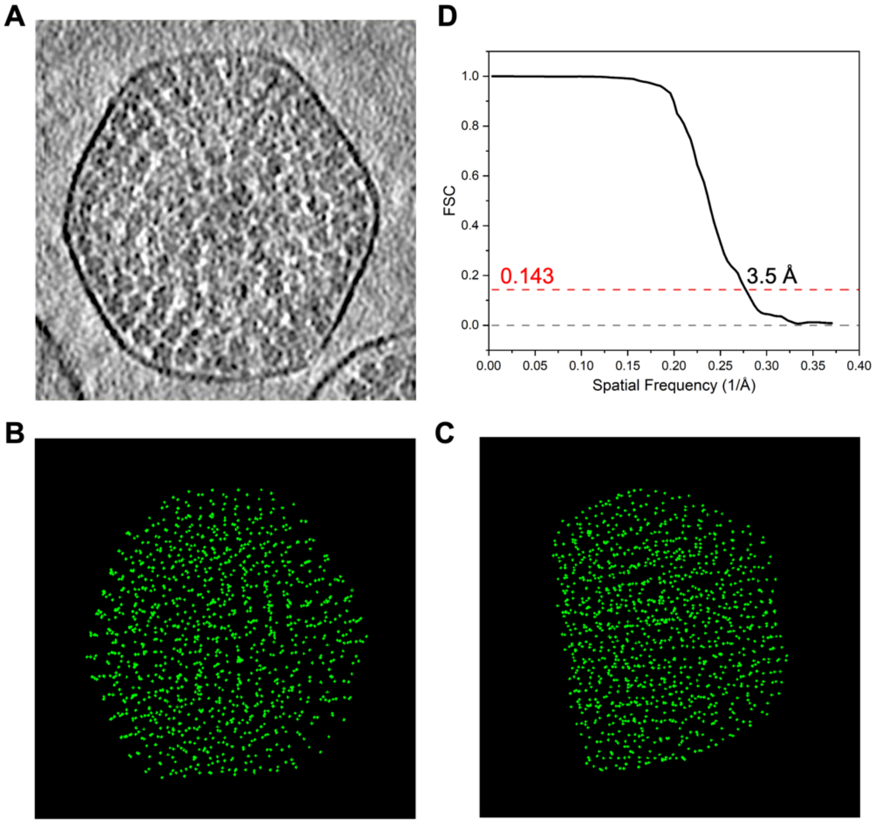


**Supplementary Figure 2. Positioning of Rubiscos in a β-carboxysome by template matching using emClarity.** (**A**) A tomographic slice of a β-carboxysome. (**B**) Top and (**C**) side projection views of template-matched Rubisco points, indicating ordered arrays of Rubiscos within the β-carboxysome. (**D**) Fourier Shell Correlation (FSC) of Rubiscos from β-carboxysomes. The resolution is 3.5 Å.at FSC of 0.143.

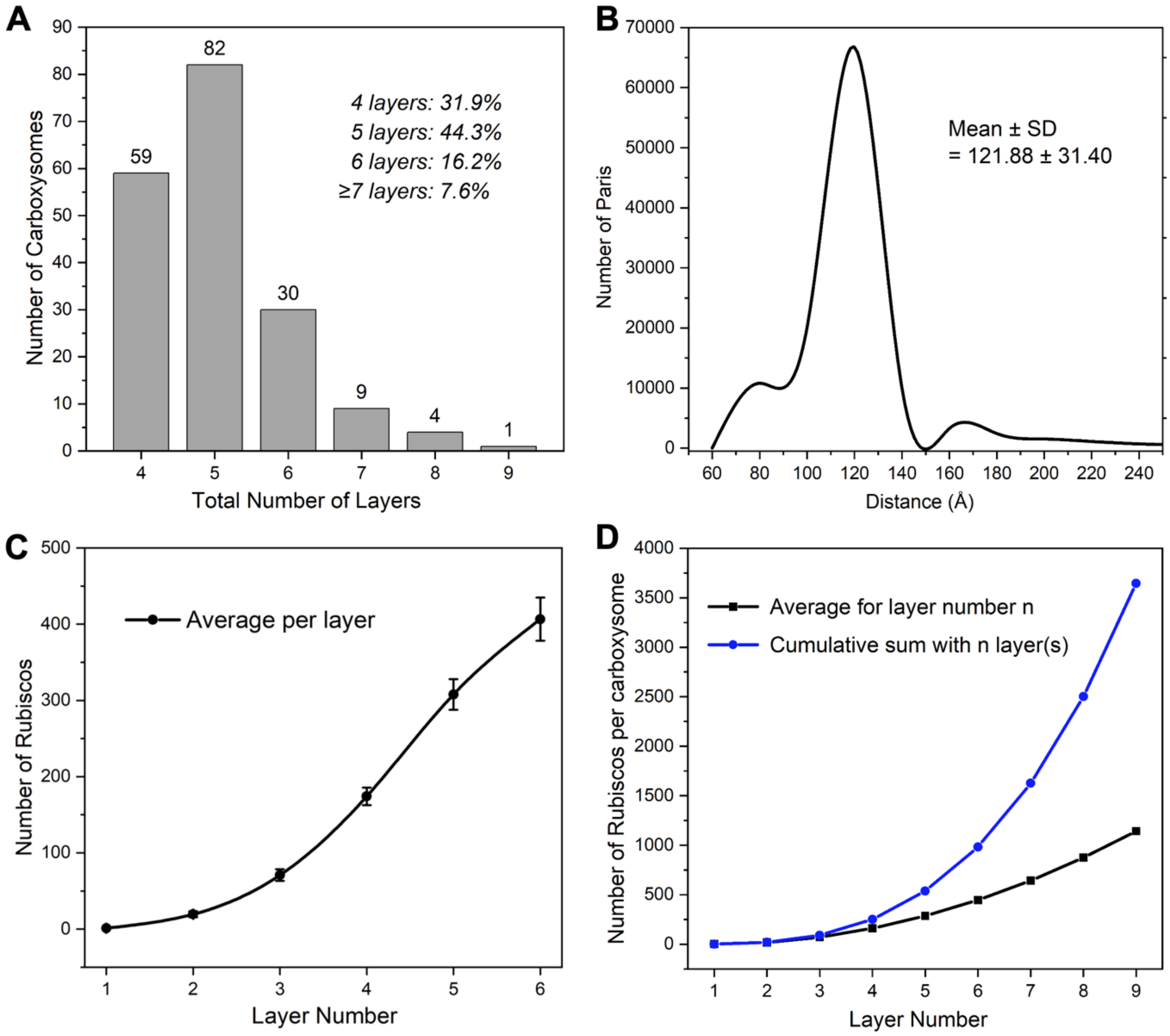


**Supplementary Figure 3. Layer profiles of Rubiscos within β-carboxysomes.** (**A**) Histogram of layer numbers in β-carboxysomes ($n$ = 185). (**B**) Pairwise Rubisco distances in β-carboxysomes ($n$, Rubisco counts = 118k). Distances within 250 Å were plotted. The average distance is 121.88 Å with a standard deviation of 31.40 Å. (**C**) Average number of Rubiscos in a β-carboxysome from layer 1 to layer 6 ($n$ = 6 carboxysome particles). (**D**) The predicted number of Rubiscos per β-carboxysome for layer number n (black) and the cumulative sum of Rubiscos in a β-carboxysome with $n$ layers (blue) ($n$ = 1-9).

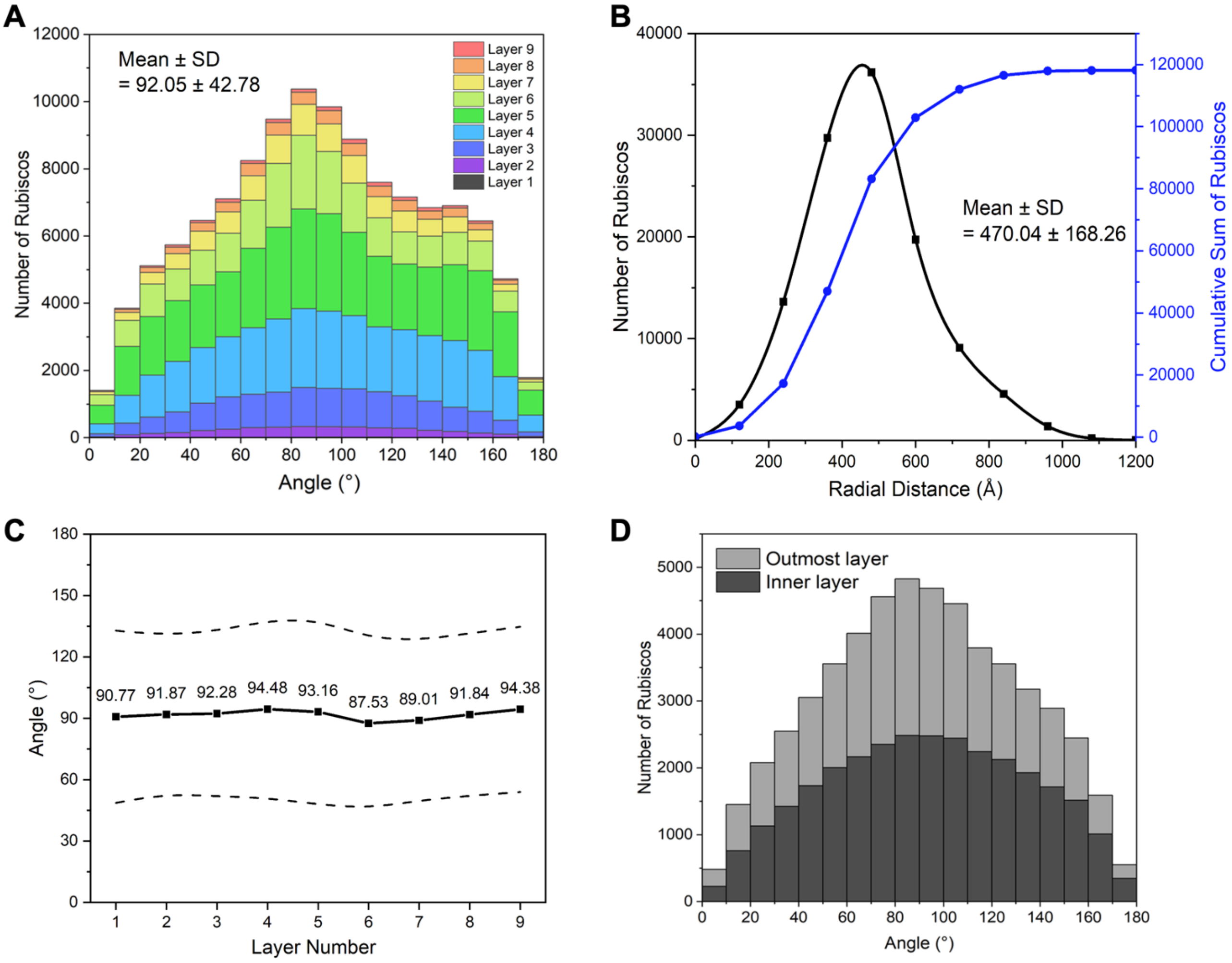


**Supplementary Figure 4. Orientation profile of Rubiscos within β-carboxysomes.** (**A-B**) Distributions of angle from radial direction (A) and radial distance (B) for Rubiscos in β-carboxysomes (*n* = 118k). (**C**) The average angle (black solid line) of Rubiscos in β-carboxysomes for each layer. The black dashed lines indicate the standard deviation of each layer (*n* = 118k). (**D**) Rubisco angle distribution comparing the outer and inner layers. Angle distribution of Rubiscos in the inner layer is shown in black and the outmost layer shown in light grey. The inner layer contains all the Rubiscos within a radial distance of 360 Å from the core, while the outmost layer has the farthest 20% Rubiscos in each β-carboxysome. The angle for the inner layer is 88.8º on average with a standard deviation of 39.7º, and the average for the outmost layer is 92.9º with a standard deviation of 41.0º.

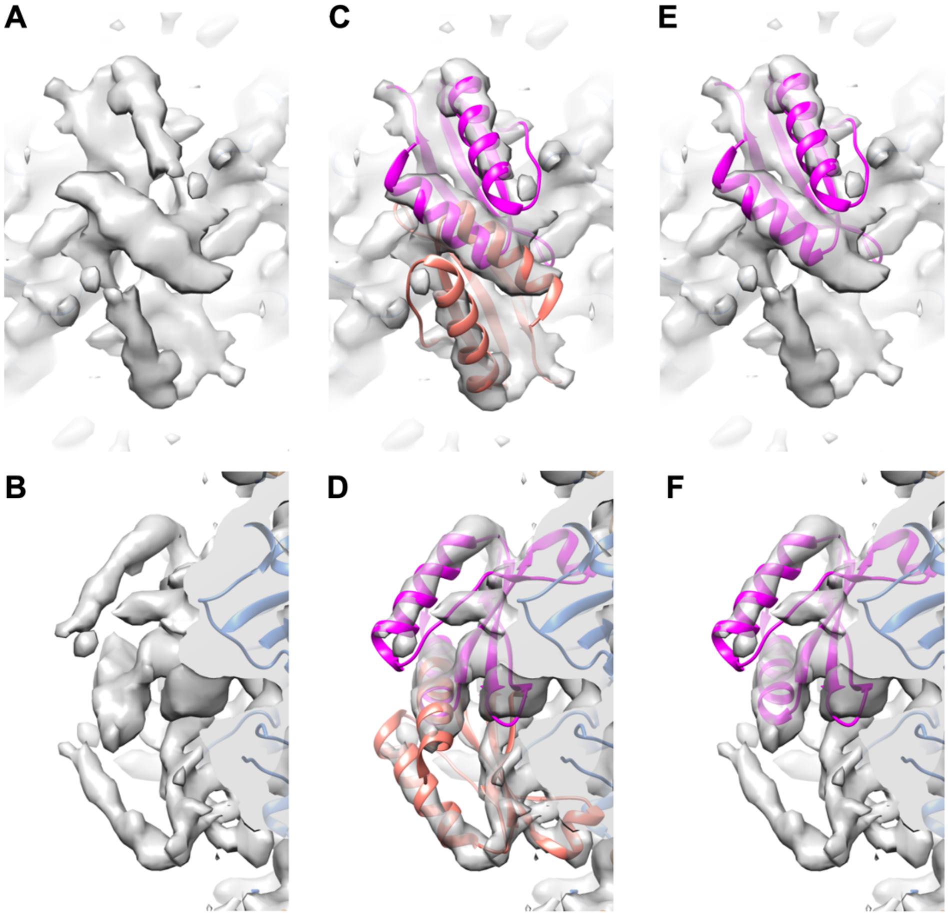


**Supplementary Figure 5. The density and fittings of CcmM SSUL domain on Rubisco. (A-B)** The top and side views of the extra density. **(C-D)** The top and side views of the extra density, fitted with atomic model of CcmM SSUL (PDB: 6HBC, magenta) to the top density. **(E-F)** The top and side views of the extra density, fitted with atomic model CcmM SSUL (PDB: 6HBC, orange) to the bottom density. D4 symmetry was applied when processing the dataset.

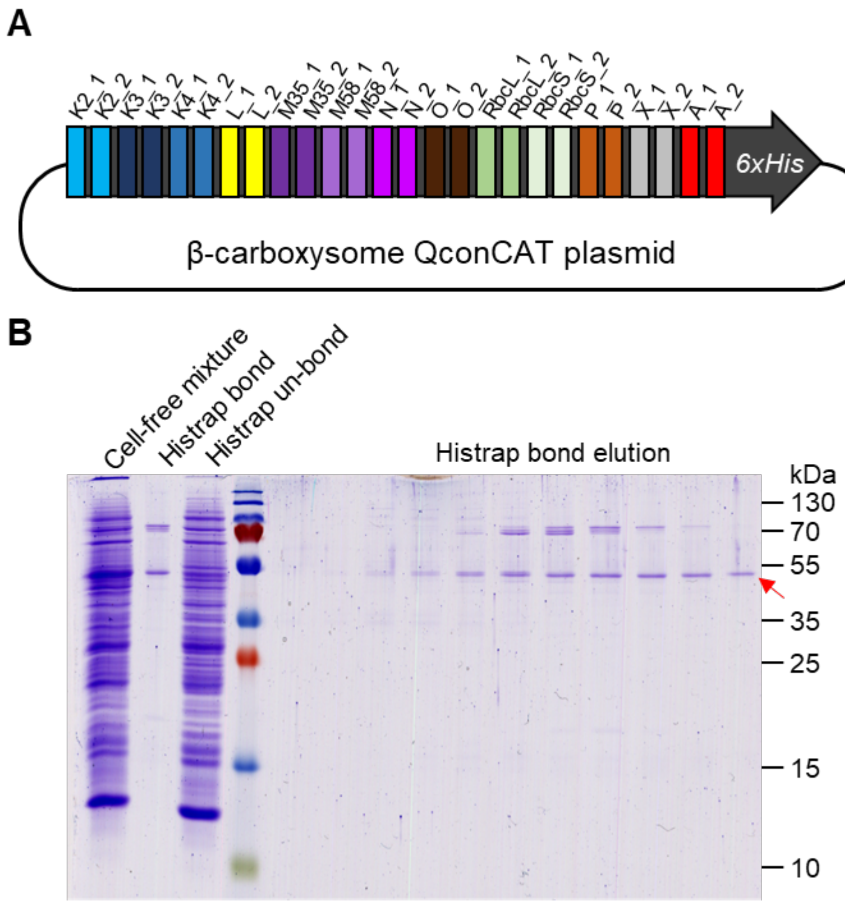


**Supplementary Figure 6. Design and preparation of QconCAT quantification.** (**A**) Design of QconCAT standard peptides, two representative peptides are selected to quantify each β-carboxysome protein. The QconCAT is also His-tagged on the C-terminus, Sequence for each peptide, full DNA and protein sequences are provided in Supplementary Tables 2 and 3; (**B**) SDS-PAGE for cell-free reaction mixture expression QconCAT peptides. The QconCAT standard at a size of 53.7 kDa was synthesized in a cell-free system and purified via the His-trap affinity column. Pure QconCAT standard was obtained in the last fraction of elution, marked by a red arrow.

**Supplementary Table 1. Cryo-ET Data collection, processing and model refinement statistics**

| Rubisco in the Syn7942 β-carboxysome, EMD-50836, PDB 9FWV | |
|---|---|
| **Data collection** | |
| Microscope | FEI Titan Krios |
| Magnification | 64k |
| Voltage | 300 kV |
| Electron dose | 120 $e^-/Å^2$ |
| Detector | Gatan K3 |
| Energy filter (slit) | Yes, 20 eV |
| Acquisition scheme | −60°/+60°, 3° dose-symmetric |
| Frame number | 10 |
| Defocus Range | −2.5 to −5.5 µm |
| Pixel Size | 1.35 Å |
| **Data processing** | |
| Symmetry imposed | D4 |
| Number of Tilt-series | 65 |
| Number of carboxysomes | 185 |
| Final particle numbers | 87,065 |
| Map resolution(Å) | 3.5 |
| FSC threshold | 0.143 |
| **Refinement** | |
| Initial Model used (PDB code) | 8BCM, 6HBC |
| Model composition | |
| Non-hydrogen atoms | 39484 |
| Residues | 4952 |
| RMSD | |
| Bond length(Å) | 0.004 |
| Bond angles(°) | 0.629 |
| MolProbity score | 2.1 |
| Clashscore | 16.41 |
| Rotamers Outliers (%) | 0.07 |
| Ramachandran Plot | |
| Favored(%) | 94.4 |
| Allowed(%) | 5.44 |
| Outliers(%) | 0.16 |

**Supplementary Table 2. QconCAT Peptides derived from tryptic proteolysis for β-carboxysome protein quantification.** The flanking sequences that recapitulate the true native primary sequence context, together with additional sequences (−3 and +3 seq) which are derived from the loop assembly synthesis of the QconCAT.

| Protein | Uniprot | Cyanobase | Candidate | −3 seq | Peptide sequence | + 3 seq | Full sequence |
|---|---|---|---|---|---|---|---|
| **CcmK2** | CCMK_SYNE7 | Synpcc7942_1421 | 1 | AAR | VTLVGYEK | IGS | AARVTLVGYEKIGS |
| **CcmK2** | CCMK_SYNE7 | Synpcc7942_1421 | 2 | SGR | VTVIVR | GDV | SGRVTVIVRGDV |
| **CcmK3** | Q31RK3_SYNE7 | Synpcc7942_0284 | 1 | SVR | GPVSEVETAVEAGLK | AVA | SVRGPVSEVETAVEAGLKAVA |
| **CcmK3** | Q31RK3_SYNE7 | Synpcc7942_0284 | 2 | AAR | VTITQYGLAESAQFFVSVR | GPV | AARVTITQYGLAESAQFFVSVRGPV |
| **CcmK4** | Q31RK2_SYNE7 | Synpcc7942_0285 | 1 | SAR | GFPPILAAADAMVK | AGR | SARGFPPILAAADAMVKAGR |
| **CcmK4** | Q31RK2_SYNE7 | Synpcc7942_0285 | 2 | SAR | FAVNIR | GDV | SARFAVNIRGDV |
| **CcmL** | CCML_SYNE7 | Synpcc7942_1422 | 1 | KVR | GTVVSTYK | EPS | KVRGTVVSTYKEPS |
| **CcmL** | CCML_SYNE7 | Synpcc7942_1422 | 2 | TYK | EPSLQGVK | FLV | TYKEPSLQGVKFLV |
| **CcmM35** | CCMM_SYNE7 | Synpcc7942_1423 | 1 | RFR | TSSWQSCAPIQSSNER | QVL | RFRTSSWQSCAPIQSSNERQVL |
| **CcmM35** | CCMM_SYNE7 | Synpcc7942_1423 | 2 | QVR | SLLNQGYR | IGT | QVRSLLNQGYRIGT |
| **CcmM58** | CCMM_SYNE7 | Synpcc7942_1423 | 1 | THK | ALIHGPAYLGDDCFVGFR | STV | THKALIHGPAYLGDDCFVGFRSTV |
| **CcmM58** | CCMM_SYNE7 | Synpcc7942_1423 | 2 | PGR | YVPSGAIITTQQQADR | LPE | PGRYVPSGAIITTQQQADRLPE |
| **CcmN** | Y1424_SYNE7 | Synpcc7942_1424 | 1 | DSR | IEIASGVCIGLGSVIHAR | GGA | DSRIEIASGVCIGLGSVIHARGGA |
| **CcmN** | Y1424_SYNE7 | Synpcc7942_1424 | 2 | AGR | SPQSSAIAHPTK | VYG | AGRSPQSSAIAHPTKVYG |
| **CcmO** | CCMO_SYNE7 | Synpcc7942_1425 | 1 | MLK | SADVTLIGYEK | TGS | MLKSADVTLIGYEKTGS |
| **CcmO** | CCMO_SYNE7 | Synpcc7942_1425 | 2 | YEK | TGSGFCTAIIR | GGY | YEKTGSGFCTAIIRGGY |
| **RbcL** | RBL_SYNE7 | Synpcc7942_1426 | 1 | FTK | DDENINSQPFQR | WRD | FTKDDENINSQPFQRWRD |
| **RbcL** | RBL_SYNE7 | Synpcc7942_1426 | 2 | WCR | DNGVLLHIHR | AMH | WCRDNGVLLHIHRAMH |
| **RbcS** | Q31NB2_SYNE7 | Synpcc7942_1427 | 1 | DCK | SPQQVLDEVR | ECR | DCKSPQQVLDEVRECR |
| **RbcS** | Q31NB2_SYNE7 | Synpcc7942_1427 | 2 | MWK | LPLFDCK | SPQ | MWKLPLFDCKSPQ |
| **CcmP** | Q31QW7_SYNE7 | Synpcc7942_0520 | 1 | AEK | AALINILQVSAIGSFGR | LFL | AEKAALINILQVSAIGSFGRLFL |
| **CcmP** | Q31QW7_SYNE7 | Synpcc7942_0520 | 2 | ALK | AAVVRPGVQFIER | LYG | ALKAAVVRPGVQFIERLYG |
| **RbcX** | Q31N04_SYNE7 | Synpcc7942_1535 | 1 | VIR | DQLAETNPAGAYR | LQV | VIRDQLAETNPAGAYRLQV |
| **RbcX** | Q31N04_SYNE7 | Synpcc7942_1535 | 2 | GLR | VMTVR | EHL | GLRVMTVREHL |
| **CcaA** | CYNT_SYNE7 | Synpcc7942_1447 | 1 | ISR | TSSDDTGIDECPVR | LPG | ISRTSSDDTGIDECPVRLPG |
| **CcaA** | CYNT_SYNE7 | Synpcc7942_1447 | 2 | MRK | LIEGLR | HFR | MRKLIEGLRHFR |

**Supplementary Table 3. DNA and protein sequences of the β-carboxysome QconCAT peptide.**

| | |
|---|---|
| Protein sequence (491 aa) | MGTRMGTREGVNDNEEGFFSARAARVTLVGYEKIGSSGRVTVIVRGDVSVRGPVSEVETAVE<br>AGLKAVAAARVTITQYGLAESAQFFVSVRGPVSARGFPPILAAADAMVKAGRSARFAVNIRG<br>DVKVRGTVVSTYKEPSTYKEPSLQGVKFLVRFRTSSWQSCAPIQSSNERQVLQVRSLLNQGY<br>RIGTTHKALIHGPAYLGDDCFVGFRSTVPGRYVPSGAIITTQQQADRLPEDSRIEIASGVCI<br>GLGSVIHARGGAAGRSPQSSAIAHPTKVYGMLKSADVTLIGYEKTGSYEKTGSGFCTAIIRG<br>GYFTKDDENINSQPFQRWRDWCRDNGVLLHIHRAMHDCKSPQQVLDEVRECRMWKLPLFDCK<br>SPQAEKAALINILQVSAIGSFGRLFLALKAAVVRPGVQFIERLYGVIRDQLAETNPAGAYRL<br>QVGLRVMTVREHLISRTSSDDTGIDECPVRLPGMRKLIEGLRHFRLAAALEHHHHHH |
| DNA sequence (1,476 bp) | ATGGGTACACGTATGGGCACCCGTGAAGGTGTTAATGATAATGAAGAAGGCTTTTTTAGCGC<br>ACGTGCAGCCCGTGTTACCCTGGTTGGTTATGAAAAAATTGGTAGCAGCGGTCGTGTTACCG<br>TTATTGTTCGTGGTGATGTTAGCGTTCGTGGTCCGGTTAGCGAAGTTGAAACCGCAGTTGAA<br>GCAGGTCTGAAAGCAGTTGCAGCCGCACGTGTTACCATTACACAGTATGGTCTGGCAGAAAG<br>CGCACAGTTTTTTGTTTCAGTGCGTGGTCCTGTTTCAGCCCGTGGTTTTCCGCCTATTCTGG<br>CAGCAGCAGATGCAATGGTTAAAGCAGGTCGTAGTGCACGTTTTGCAGTTAATATTCGCGGT<br>GATGTGAAAGTTCGTGGCACCGTTGTTAGCACCTATAAAGAACCGAGTACATACAAAGAACC<br>GTCACTGCAGGGTGTTAAATTTCTGGTTCGTTTTCGTACCAGCAGCTGGCAGAGCTGTGCAC<br>CGATTCAGAGCAGCAATGAACGTCAGGTTCTGCAGGTTCGTAGCCTGCTGAATCAGGGTTAT<br>CGTATTGGTACAACCCATAAAGCACTGATTCATGGTCCGGCATATCTGGGTGATGATTGTTT<br>TGTTGGTTTTCGTAGTACCGTTCCGGGTCGTTATGTTCCGAGCGGTGCAATTATTACCACAC<br>AGCAGCAGGCCGATCGTCTGCCGGAAGATAGCCGTATTGAAATTGCAAGCGGTGTTTGTATT<br>GGTCTGGGTAGCGTTATTCATGCACGTGGTGGTGCAGCCGGTCGTAGTCCGCAGAGCAGCGC<br>AATTGCACATCCGACCAAAGTTTATGGTATGCTGAAAAGCGCAGATGTGACCCTGATCGGCT<br>ATGAGAAAACCGGTAGTTACGAAAAAACCGGCAGCGGTTTTTGTACCGCAATTATTCGTGGT<br>GGCTATTTCACCAAAGATGACGAAAACATTAATAGCCAGCCGTTTCAGCGTTGGCGTGATTG<br>GTGTCGTGATAATGGTGTTCTGCTGCATATTCATCGTGCAATGCATGATTGTAAAAGTCCGC<br>AGCAGGTTCTGGATGAAGTTCGTGAATGTCGTATGTGGAAACTGCCGCTGTTTGATTGCAAA<br>TCACCGCAGGCAGAAAAAGCAGCCCTGATTAATATCCTGCAGGTTAGCGCCATTGGTAGCTT<br>TGGTCGTCTGTTTCTGGCACTGAAAGCCGCAGTGGTTCGTCCGGGTGTTCAGTTTATTGAAC<br>GTCTGTATGGTGTTATTCGTGATCAGCTGGCCGAAACCAATCCGGCAGGCGCATATCGTCTG<br>CAGGTAGGTCTGCGTGTTATGACCGTTCGTGAACATCTGATTAGCCGTACCAGCTCAGATGA<br>TACCGGCATTGATGAATGTCCGGTTCGTCTGCCTGGTATGCGTAAACTGATTGAAGGTCTGC<br>GCCATTTTCGTCTGGCAGCGGCACTGGAACATCATCACCATCATCATTAG |

**Supplementary Table 4. Comparison of the protein ratios of β-carboxysomes from Syn7942 and α-carboxysomes from *H. neapolitanus* (Sun *et al*., 2022).** *Carbonic anhydrases are considered as the monomeric forms for comparison.

| | Ratio | |
|---|---|---|
| | **Syn7942 β-CB** | ***H. neap* α-CB** |
| Rubisco:scaffolding protein (CcmM or CsoS2) | 1:1.2 | 1:1 |
| Rubisco:carbonic anhydrases* | 33:1 | 31:1 |
| Scaffolding short:long isoforms (CcmM35/58 or CsoS2A/B) | 6.3:1 | 1.3:1 |
| Major shell hexamer:shell-interacting protein (CcmN or CsoS2B) | 247:1 | 5.1:1 |
| Major:minor shell hexamers | 86:1 | 8:1 |
| Shell proteins:cargos | 2:1 | 2:1 |
| Hexamers:trimers | 145:1 | 336:1 |
| Hexamers:pentamers | 347:1 | 134:1 |
